# Microglia determine an immune-challenged environment and facilitate ibuprofen action in human retinal organoids

**DOI:** 10.1101/2024.09.20.614136

**Authors:** Verena Schmied, Medina Korkut-Demirbaş, Alessandro Venturino, Sandra Siegert

## Abstract

Prenatal immune challenges pose significant risks to human embryonic brain and eye development. However, we still lack knowledge about the safe usage of anti-inflammatory drugs during pregnancy. While human induced pluripotent stem cells (hIPSC)-derived brain organoid models have started to explore functional consequences upon viral stimulation, these models commonly lack microglia. Yet, microglia are susceptible to and promote inflammation and will influence the effects.

Here, we generate hIPSC-derived microglia precursor cells and assemble them into retinal organoids to dissect their interplay with the developing nervous system and expose them to the immunostimulant poly(I:C). We first identified successful hIPSC-derived microglia (iMG) integration into the retinal organoid at the time when the outer plexiform layer forms. To improve ganglion cell surveillance, we adapted the retinal organoid model and found that the ganglion cell number significantly decreased only with iMG presence. While poly(I:C) exposure alters the iMG phenotype, it does not hinder their interaction with ganglion cells. Furthermore, iMG significantly enhances the supernatant’s inflammatory secretome and increases retinal cell proliferation. Simultaneous exposure with the non-steroidal anti-inflammatory drug (NSAID) ibuprofen dampens the poly(I:C)-mediated changes of the iMG phenotype and ameliorates cell proliferation. Remarkably, while poly(I:C) disrupts neuronal calcium dynamics independent of iMG, ibuprofen rescues this effect only if iMG are present. This effect depends on cyclooxygenase 1 in microglia and cyclooxygenase 2, common NSAID targets.

These findings underscore the importance of microglia in the context of prenatal immune challenges and provide insight into the mechanisms by which ibuprofen exerts its protective effects during embryonic development.

## Introduction

Prenatal exposure to infections can be detrimental to human embryonic development [1, 2]. Certain infectious diseases like rubella belonging to the TORCH complex (Toxoplasmosis, Others, Rubella, Cytomegalovirus, Herpes) can be vertically transmitted from pregnant women to their fetus, resulting in malformations of the fetal brain and eye [3–6]. Medication is recommended to a certain degree to treat inflammatory symptoms during pregnancy, but there are significant knowledge gaps on the effects of anti-inflammatory drugs on embryonic development [7].

Brain organoids derived from human induced pluripotent stem cells (hIPSCs) provide a unique strategy to investigate the consequences of prenatal inflammation, which we refer to as neuro-immune challenge, and drug exposure on neuronal development. Specifically, retinal organoids are one of the first established brain region-specific models [8], in which the developmental trajectories and cytoarchitecture are well-defined [9, 10] and match anatomical observations in human fetal retinal development, like the formation of the ganglion cell layer and the outer plexiform layer (OPL) [11]. At the same time, neuroectodermal-derived organoids commonly lack mesodermal-derived brain-resident macrophages [12], which colonize the human fetal brain and eye between gestation week (GW) 4.5 and 5 [13, 14]. Once in the neuronal environment, these microglia have multifunctional developmental tasks demonstrated in the rodent nervous system. They regulate, amongst others, the number of neural precursor cells [15, 16], axonal outgrowth and neuronal wiring [17] as well as synaptogenesis and pruning [18, 19] across various brain regions [20–22] [20–23]. Microglia maintenance and survival depend on the colony-stimulating factor 1 receptor (CSF1R) [24–26], which, when inhibited, affects the total number of neurons and the macroglia cell populations consisting of astrocytes, oligodendrocytes and also their migration, distribution, and the functional connectivity [17, 20, 27–30]. Embryonic death and brain malformation have been reported in humans harboring homozygous mutations within the CSF1R genome [31, 32].

In recent years, protocols have been developed to generate hIPSC-derived microglia precursor cells (preMG), which acquire microglia-like cell (iMG) properties once integrated into neuroectodermal tissue and exposed to the environmental cues [33, 33–37, 37, 38]. Latest studies demonstrate that iMG promote brain organoid maturation [38] and fine-tune their neuronal environment at the cellular and synaptic levels [39–41]. Thus, microglia integration seems relevant to mimic *in vivo* human brain development.

Human cerebral organoids have been used to model the consequences of TORCH viruses to which Zika also belongs, and found reduced neuronal progenitors [42–46]. However, due to the lack of microglia, these studies are limited in their insights into the inflammatory response and its consequences on human embryonic development. Microglia are susceptible to environmental cues beyond pathogens [47], including inflammatory mediators such as cytokines and chemokines [48–50]. In rodent models, prenatal neuro-immune challenges induce microglia to express receptors to sense pathogens and inflammatory mediators [51] and affect microglia properties such as morphology, motility, and their actual number [16, 17, 52–54]. In parallel, these immune challenges also affect neurogenesis [15, 16, 22], neuronal differentiation [55], synaptogenesis [56, 57], and synaptic pruning [58, 59], which are tasks that microglia are involved in regular human brain development.

Here, we determined the consequences of microglia exposure to a prenatal neuro-immune challenged environment and subsequent treatment with the non-steroidal anti-inflammatory drug (NSAID) ibuprofen in microglia-assembled retinal organoids (iMG-RO).

First, we identified the optimal timepoint to investigate microglia-neuron interaction in the hIPSC-derived 3D-retinal organoid (RO) that mimics the regular retinal development the closest [60]. Then, we developed a 2D model system (_diss_RO) based on the same timeline to counteract the previously reported ganglion cell loss in RO [10, 61, 62] and to avoid known challenges of organoid-to-organoid variability in size, shape, and cell type composition [10, 63, 64], as well as diffusion biases of drugs. In this iMG-_diss_RO model, we observed that iMG actively interact with and phagocytose retinal ganglion cells.

Next, we modeled a prenatal neuro-immune challenge by exposing the culture to the immunostimulant *polyinosinic: polycytidylic acid*. This poly(I:C) mimics a viral-mediated response and activates the toll-like receptor 3 (TLR3) [65]. TLR3 stimulation induces a downstream signaling cascade involving NFkB- and interferon pathways, resulting in cytokines and chemokines release [47, 66, 67]. Furthermore, poly(I:C) directly acts on microglia as they upregulate *TLR3* mRNA expression [68]. We investigated the consequences of poly(I:C)-mediated immune challenge in our iMG-_diss_RO model and identified a microglia-dependent inflammatory signature and increased retinal cell proliferation. To evaluate the effects of anti-inflammatory drugs on the identified consequences, we focused on the NSAID ibuprofen, which can be taken cautiously during the first half of the pregnancy [69]. Ibuprofen targets cyclooxygenase 1 and 2 (*PTGS1*/COX1, *PTGS2*/COX2, respectively) and therefore prevents the conversion of arachidonic acid into prostaglandins like PGE2 [70, 71]. In the presence of ibuprofen, poly(I:C)-mediated effects on microglia were dampened, and the neuronal phenotypes were restored. Yet, this beneficial effect depended on the selective role of PTGS1 expressed by iMG since ibuprofen did not show this rescue in cultures without iMG.

Our study highlights the interplay of microglia with neurons under prenatal neuro-immune challenges and the consequences of drug treatments. In light of future drug testing and known species-specific differences in the induction of microglia inflammatory response [72, 73], our results emphasize the relevance of including iMG into neuronal organoids.

## Results

### OPL formation aligns with successful iMG integration into retinal organoids

To generate retinal organoids (RO), we differentiated the human induced pluripotent stem cell (hIPSC) line F49B7, which has been recently analyzed for its transcriptional cell diversity across different time points of RO differentiation [10]. We monitored the retinal cup formation under brightfield microscopy over 30 weeks (**Figure 1a**).

**Figure 1-.**
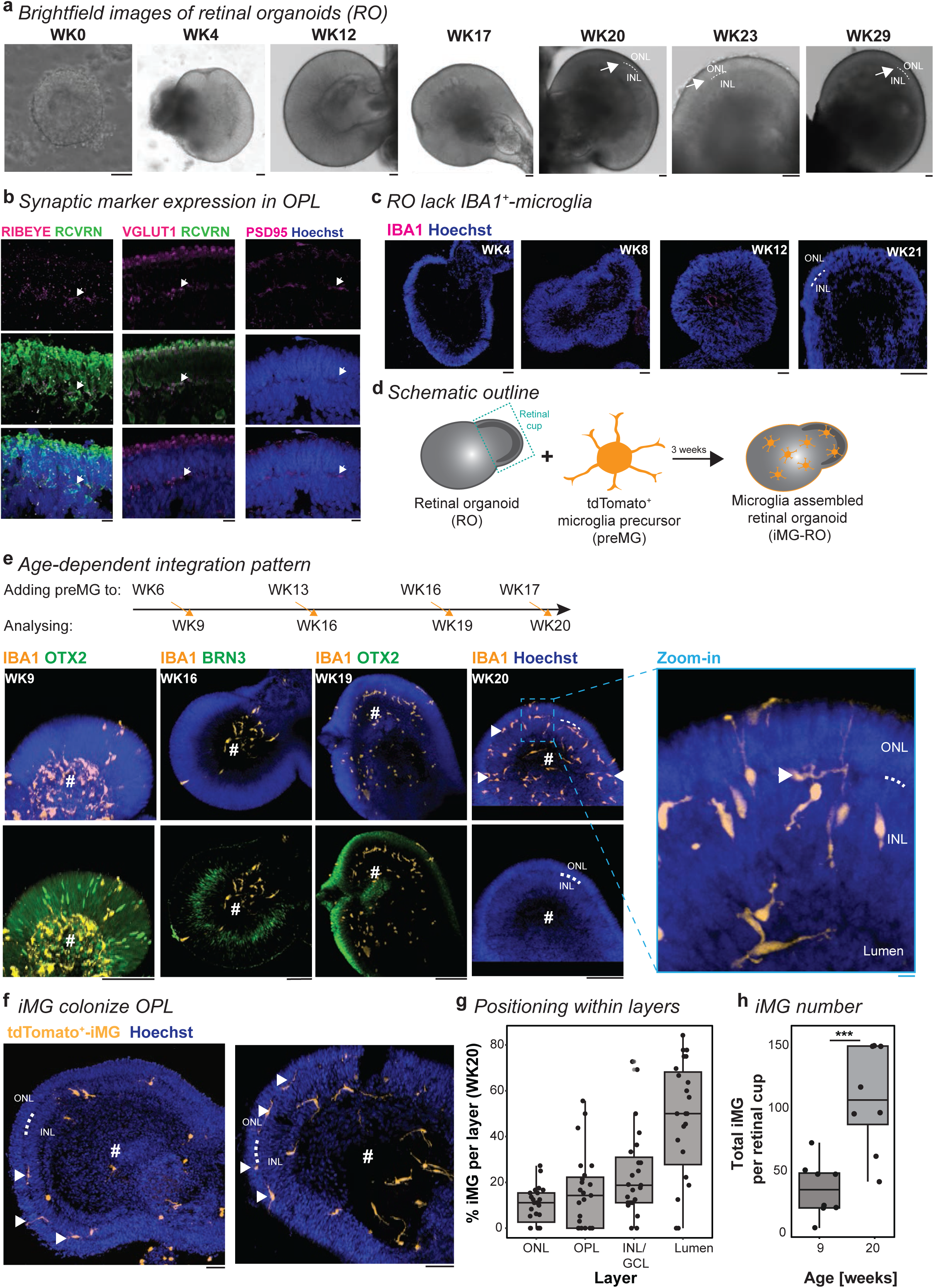
Microglia colonize retinal layers after OPL formation. **a**, Representative brightfield images focusing on the retinal cup at selected RO differentiation time points. Arrow and dashed line: outer plexiform layer formation, visible from WK20 onwards. Scale bar: 100 µm. **b**-**c**, Images of RO cryostat sections counterstained with the nuclei-dye Hoechst (blue) and immunostained for **b**, photoreceptors with RCVRN (green) and the ribbon synapse marker RIBEYE (magenta, left); presynaptic marker VGLUT1 (magenta, middle); and postsynaptic marker PSD95 (magenta, right) at WK20. White arrow: outer plexiform layer. Scale bar: 10 µm. **c**, IBA1 (magenta) at WK4, WK8, WK12, and WK21. Scale bar: 100 µm. **d**, Experimental schematic to generate iMG-RO. **e**, Maximum intensity projection image of entire iMG-RO counterstained with the nuclei-dye Hoechst (blue) at different time points of preMG application and collection of iMG-RO as outlined in the schematic. Immunostaining for IBA1 (orange), BRN3 (green, for WK16), and OTX2 (green, for WK9 and WK19). White arrowhead: iMG positioned in OPL. White dashed line: OPL. #: retinal cup lumen. Scale bar: 100 µm. Zoom-in: 10 µm. **f**, Images of iMG-RO cryostat sections counterstained with the nuclei-dye Hoechst (blue) and tdTomato^+^-iMG (orange). White arrowhead: iMG located in OPL. White dashed line: OPL. #: retinal cup lumen. Scale bar: 50 µm. **g**-**h**, Boxplot. **g**, Percent of iMG in the ONL, OPL, INL, GCL, and within the retinal cup lumen (#) at WK20. Each dot: one cryostat section of an independent retinal cup. **h**, Total number of iMG integrated per retinal cup at WK9 and WK20. Each dot represents an entire retinal cup. Students’s t-test. *** p<0.001. For detailed statistical analysis, see **Supplementary Table 4**. Abbreviations: BRN3: brain-specific homeobox/POU domain protein 3B. GCL: ganglion cell layer. hIPSC: human induced pluripotent stem cell. IBA1: ionized calcium-binding adapter molecule 1 (alternative name: AIF1). INL: inner nuclear layer. iMG: microglia-like cell. iMG-RO: microglia assembled 3D-retinal organoid. ONL: outer nuclear layer. OPL: outer plexiform layer. OTX2: orthodenticle homeobox 2. preMG: microglia precursor cells. PSD95: postsynaptic density protein 95. RCVRN: recoverin. RO: 3D-retinal organoid. VGLUT1: vesicular glutamate transporter 1. WK: week after the start of RO differentiation.

Once the outer plexiform layer (OPL) was visible around week (WK) 20, we performed immunostaining of the RO for synaptic markers (**Figure 1b**). The OPL showed expression of the presynaptic markers RIBEYE and VGLUT1 and the post-synaptic marker PSD95.

Furthermore, we confirmed the existence and the location of the different cell types within their expected nuclear layer at WK20, such as RCVRN^+^-/ OTX2^+^-/ CALB2^+^-photoreceptors in the outer nuclear layer and OTX2^+^/ CALB2^+^-bipolar cells, CALB2^+^-amacrine cells, CALB1^+^-horizontal- and amacrine cells, and CHAT^+^-amacrine cells in the inner nuclear layer (**Supplementary Figure 1a**). Few BRN3^+^-ganglion cells localized close to the RO lumen.

RLBP1^+^-Müller glial cells expanded their processes across all layers. OPL formation and cell type expression patterns matched the anticipated timeline observed in human fetal tissue studies [74–76].

Human microglia have been shown to accumulate at the optic disc between GW10-13 and then populate the OPL between GW20-25 [60]. When we stained the RO for the microglia-associated marker IBA1 [77], we did not find innately developing IBA1^+^-microglia within the retinal cup at any collected time points (**Figure 1c**). This is in line with our previous observations [34] and confirms the sequencing data at weeks 30 and 38 by Cowan *et al.*, which failed to identify microglia signature gene transcripts like IBA1/AIF1, CX3CR1, PU.1/SPI1, and P2RY12 (**Supplementary Figure 1b**) [78–81]. Therefore, we focused on a microglia-assembled retinal organoid (iMG-RO) model, for which we developed a hIPSC line expressing the red fluorescent protein from the AAVS1 locus (**Supplementary Figure 2a-c**) [82]. First, we confirmed that the hIPSC line remained pluripotent (**Supplementary Figure 2d**) and successfully differentiated into tdTomato^+^/IBA1^+^-microglia precursor cells (preMG) expressing the previously described and expected preMG-markers [34] (**Supplementary Figures 2e-k**). Then, we added tdTomato^+^-preMG to ROs at WK 6, 13, 16, or 17 of RO differentiation and followed their integration (**Figure 1d-e**). Independent of the differentiation week of the organoid, tdTomato^+^-preMG attached to the developing outer nuclear layer within 24h (**Supplementary Figure 3a**). After a few days, the initially roundly-shaped preMG infiltrated into the RO and adapted their morphology into a bipolar profile, which spanned throughout the layers projecting towards the lumen of the retinal cup (**Supplementary Figure 3b**). Differences in the preMG integration pattern correlated with the OPL formation. Before WK20, iMG preferentially accumulated in the lumen close to BRN3^+^-ganglion cells and rarely colonized the developing nuclear layers (**Figure 1e**). After WK20, iMG integrated fully into the OPL, or they extended their processes toward it (**Figure 1e-f**). The position of the iMG soma indicated a spatial distribution across all retinal layers (**Figure 1g**), and the total number of iMG significantly increased from WK9 to WK20 (**Figure 1h**). Overall, we determined WK20 as the time point, which aligns with the microglia integration and spatial distribution pattern in human retinal development [60].

### iMG control ganglion cell number in adapted 2D-RO model with improved ganglion cell survival

In human fetal tissue, the ganglion cell layer fully forms by GW24 [11], and its formation is accompanied by extensive cell loss peaking between GW16 and 21 [83]. Microglia have been shown to interact with newborn BRN3^+^-ganglion cells and reduce their density in the rodent retina [20]. To recapitulate this phenotype in human RO is challenging due to the reported and also in our hands verified gradual loss of retinal ganglion cells with increasing maturation (**Figure 2a-b**) [10, 61, 62]. Therefore, we adapted recent protocols that dissociate 3D organoids, plated them as 2D cultures, and validated cortical network activity reestablishment [84, 85]. Dissociated retinal organoid culture (_diss_RO) will allow us to minimize diffusion biases and compare treatment paradigms directly because the wells derive from the same pool of dissociated WK15 ROs, circumventing organoid-to-organoid variability. Until WK20, retinal cells will have had sufficient time to successfully reform their synaptic connections [86]. First, we compared the cell type composition and density to the age-matched ROs (**Figure 2c-d**). We found that the percentage of each cell type was similar between RO and _diss_RO with the exceptions of CALB1 and BRN3, which both significantly increased in _diss_RO (**Figure 2d**). Importantly, brain-derived neurotrophic factor (BDNF) in the culture medium supported BRN3^+^-ganglion cell survival in _diss_RO compared to RO (**Supplementary Figure 4a**).

**Figure 2-.**
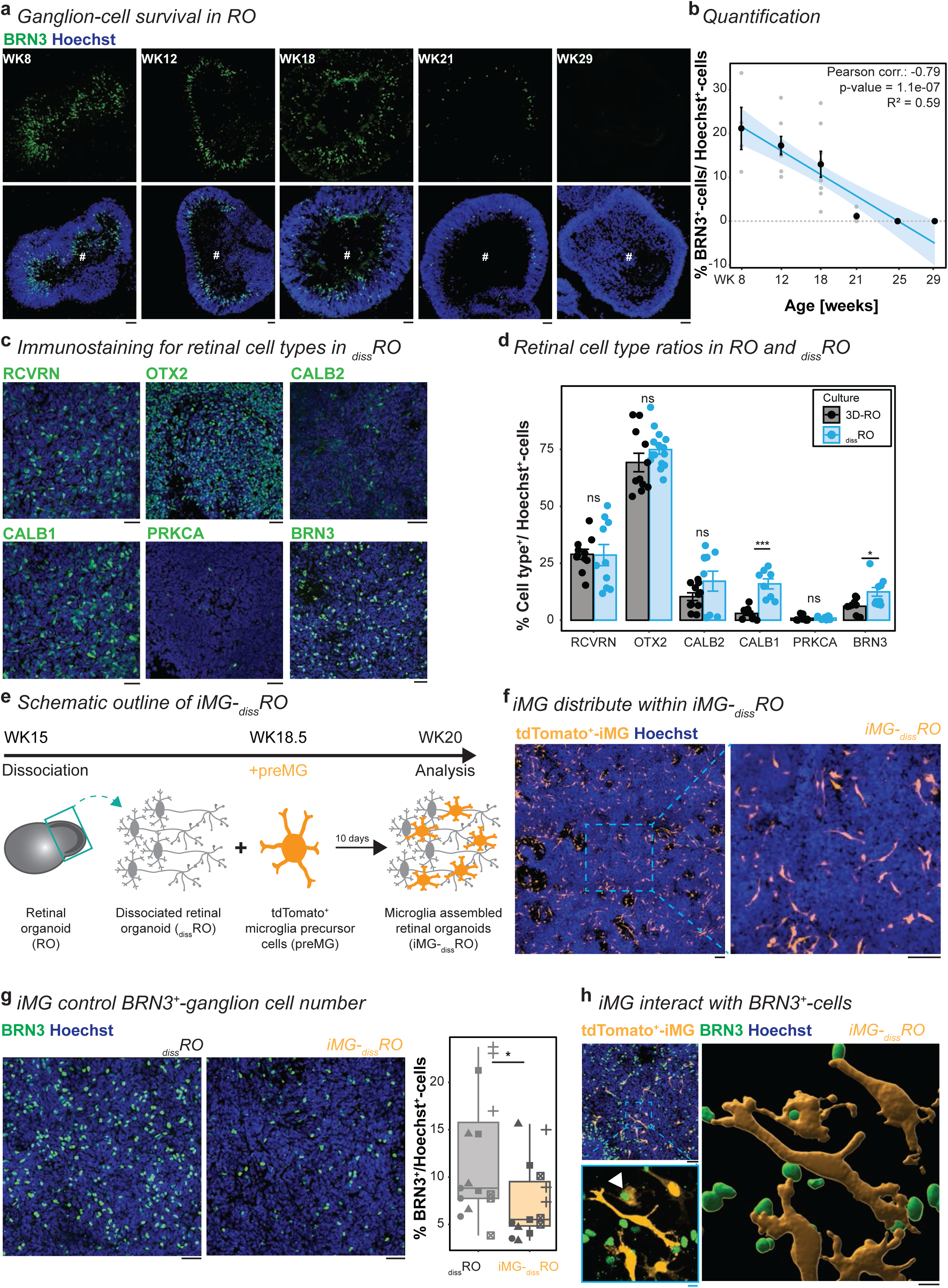
iMG interact with BRN3^+^-ganglion cells in dissociated retinal organoids. **a-b**, Gradual ganglion cell loss with RO maturation. **a**, Images of RO cryostat sections counterstained with the nuclei-dye Hoechst (blue) collected at WK8, 12, 18, 21, 29 and immunostaining for BRN3 (green). Scale bar: 50 µm. **b**, Scatterplot of BRN3^+^-cells relative to Hoechst^+^-cells per cryostat section with SEM and trend curve. Pearson correlation with a significant negative correlation between the differentiation age and the BRN3^+^-cells number. **c-d**, Retinal cell types in _diss_RO at WK20. Retinal cell type markers: RCVRN for photoreceptors, OTX2 for photoreceptors and bipolar cells, CALB2 for photoreceptors, bipolar- and amacrine cells, CALB1 for amacrine-, horizontal cells, PRKCA for rod bipolar cells, and BRN3 for ganglion cells. **c**, Immunostaining for retinal markers (green) and nuclei-dye Hoechst (blue). Scale bar: 50 µm. **d**, Bar chart with SEM of retinal cell types relative to Hoechst^+^-cells in ROs (black) and _diss_RO (blue). Each dot: cryostat section of individual ROs (black) or a field of view in _diss_RO (blue). Student’s t-test except for CALB1 (Wilcoxon rank-sum test). **e**, Experimental timeline to generate iMG-_diss_RO. At WK15, retinal cups dissociated and plated as _diss_RO. At WK18.5, independently differentiated tdTomato^+^-preMG were added. iMG-_diss_RO was analyzed ten days later at WK20. **f**, Image of iMG distribution within iMG-_diss_RO (orange) at WK20, counterstained with the nuclei-dye Hoechst (blue). Scale bar: 100 µm. **g**, Image of _diss_RO (left) and iMG-_diss_RO (right) at WK20 counterstained with the nuclei-dye Hoechst (blue) and immunostained for BRN3 (green). Scale bar: 50 µm. Next, boxplot of BRN3^+^-ganglion cells relative to Hoechst^+^-cells in _diss_RO (grey) and iMG-_diss_RO (orange). Symbols: single ROI of three biological replicates from five independent differentiation. Wilcoxon rank-sum test. **h**, Representative image of BRN3^+^-ganglion cells (green), tdTomato^+^-iMG (orange), and the Hoechst (blue) of iMG-_diss_RO at WK20. Scale bar: 50 µm. Zoom-in: 3D-surface rendering of a region of interest. White arrowhead: iMG engulfing BRN3^+^-cell. Scale bar: 10 µm. For detailed statistical analysis, see **Supplementary Table 4**. ***p < 0.001. *p < 0.05. ^ns^p > 0.05, not significant. Abbreviations: BRN3: brain-specific homeobox/POU domain protein 3B. CALB1: calbindin. CALB2: calretinin. iMG-_diss_RO: microglia-assembled dissociated retinal organoids. iMG-RO: microglia assembled 3D-retinal organoid. iMG: microglia-like cells. preMG: microglia precursor. PRKCA: protein kinase C alpha. OTX2: orthodenticle homeobox 2. RO: 3D-retinal organoid. _diss_RO: dissociated retinal organoid cultures without iMG. RCVRN: recoverin. ROI: region of interest. SEM: standard error of the mean. WK: week after the start of RO differentiation.

Next, we added tdTomato^+^-preMG to _diss_RO at WK18.5 (iMG-_diss_RO, **Figure 2e**). After ten days in culture, iMG were distributed across the plate, representing 2.61% ±1.13 of the total Hoechst^+^-nuclei number (**Figure 2f**) and expressed the expected iMG-associated markers (**Supplementary Figure 4b**). iMG removed cellular debris exemplified in their processes surrounding fragments labeled with the apoptotic marker cleaved caspase-3 (CCAS3, **Supplementary Figure 4c**). iMG-_diss_RO also contained fewer Hoechst^+^-nuclear fragments than cultures without iMG (**Supplementary Figure 4d-e**). Similar to rodent studies [20], the number of BRN3^+^-ganglion cells significantly reduced in iMG-_diss_RO compared to _diss_RO (**Figure 2g**). Furthermore, 22.53% ± 7.13 % of all BRN3^+^-ganglion cells positioned within a 5 μm radius of iMG and an average of 1.72 ± 1.70 iMG engulfed BRN3^+^-ganglion cell body (**Figure 2h**), indicating their role in regulating neuron number during development.

### Poly(I:C) affects iMG phenotype without interfering with the ganglion cell interaction

To mimic a prenatal neuro-immune challenge in our WK20 culture, we applied poly(I:C) for 24h (**Figure 3a**), which activates a TLR3 response cascade triggering downstream signaling pathways related to immune defense [87, 88]. We confirmed that in preMG culture, *TLR3* mRNA level significantly increased after poly(I:C) stimulation (**Figure 3b**), supporting a direct effect of poly(I:C). Next, we monitored iMG activity for 20 minutes in iMG-_diss_RO (**Supplementary video 1-2**). We found that iMG surveillance significantly increased compared to the control condition without poly(I:C) (**Figure 3c**). Furthermore, iMG enlarged their surface area (**Figure 3d-e**), a phenotype commonly found in reactive microglia [89, 90]. Based on these iMG phenotypes, we revisited the previously observed iMG-ganglion cell interaction (**Figure 2g-h**). iMG engulfed a comparable number of ganglion cells to age-matched, untreated control condition (**Figure 3f**). When we analyzed the number of BRN3^+^-ganglion cells, we observed a trend towards an increase in poly(I:C)-treated culture, but this effect was insignificant (**Figure 3g**). We thus also investigated whether the iMG interaction with BRN3^+^-ganglion cells is altered and determined the iMG position within a 5 μm radius of BRN3^+^-labeling. The proximity measurement did not reveal an apparent difference between poly(I:C)-stimulated and non-stimulated conditions (**Figure 3h-i**), suggesting that poly(I:C) does not have an immediate effect on the iMG developmental task to regulate the ganglion cell number.

**Figure 3-.**
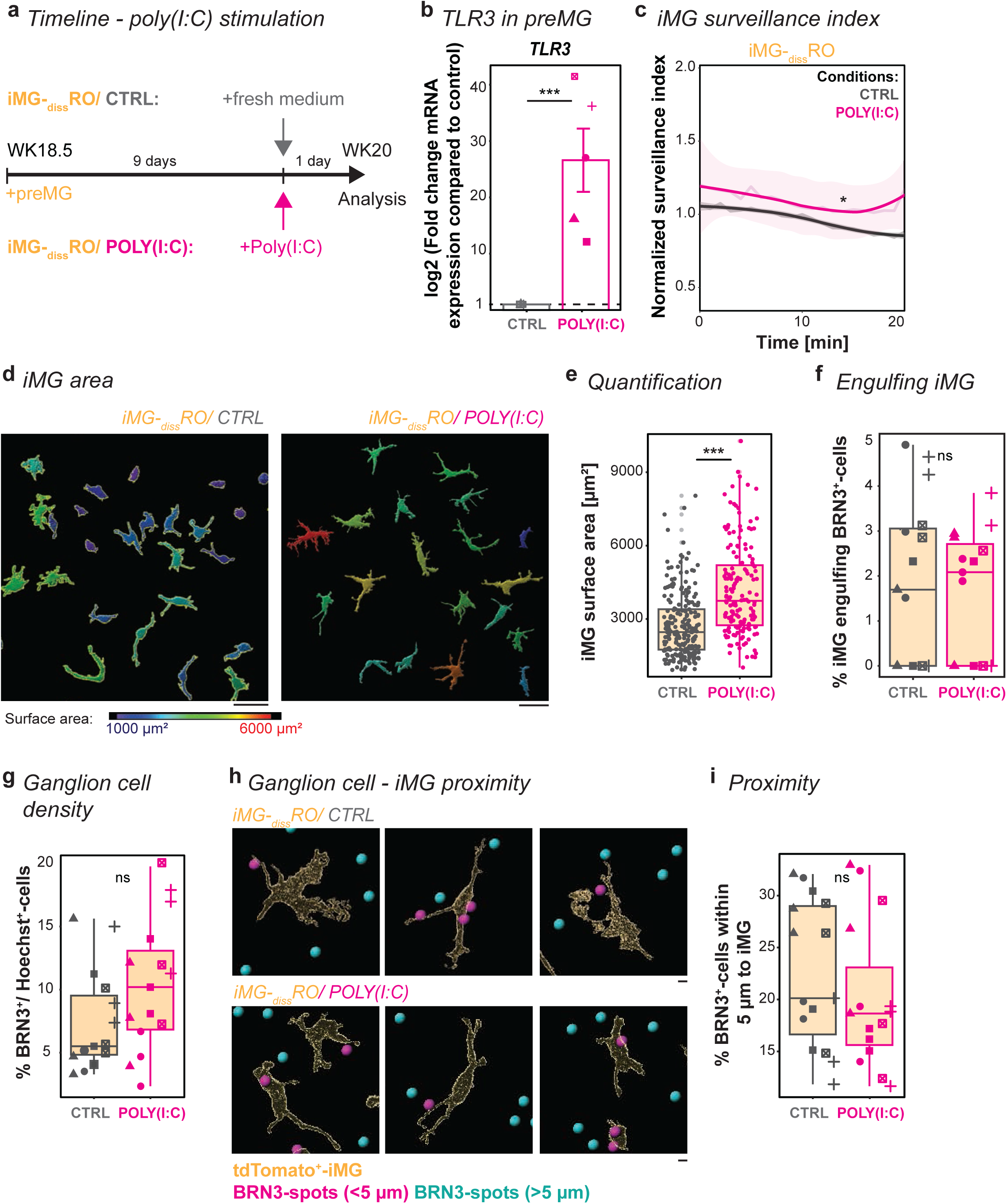
iMG respond to poly(I:C) stimulation but still interact with ganglion cells. **a**, Experimental timeline. At WK18.5, preMG are added to _diss_RO. After nine days, the culture medium was replaced with fresh medium either containing poly(I:C) (magenta) or without as a control (CTRL, grey). Analysis was performed 24 hours later on day 10. **b**, RT-qPCR for TLR3 of preMG after CTRL or poly(I:C) stimulation. Bar chart with SEM: Mean mRNA transcript log2-fold changes compared to CTRL. Symbol: mean of technical triplicate from five independent differentiations. One sample t-test. **c**, iMG-_diss_RO live imaging for 20 minutes after 24 hours of CTRL or poly(I:C) stimulation. iMG surveillance index normalized to the mean surveillance of the cells in CTRL with a 95% confidence interval. Four independent differentiations. Wilcoxon rank-sum test. **d-e**, iMG surface area quantification. **d**, iMG surface rendering for CTRL (left) and poly(I:C) (right), color-coded based on surface area: blue = 1000 µm^2^ to red = 6000 µm^2^. Scale bar: 50 µm. **e**, Boxplot of individual iMG surface areas in iMG-_diss_RO for CTRL and poly(I:C). iMG were collected from five independent differentiations. Wilcoxon rank-sum test. **f**, Boxplot quantifying the number of iMG engulfing BRN3^+^-cells in iMG-_diss_RO for CTRL and poly(I:C). Symbols: single ROI of three biological replicates from five independent differentiations. Wilcoxon rank-sum test. **g**, Boxplot determines ganglion cell density based on BRN3^+^-cells relative to Hoechst^+^-cells in iMG-_diss_RO for CTRL and poly(I:C). Symbols: single ROI of three biological replicates from five independent differentiations. Wilcoxon rank-sum test. **h-i**, Ganglion cell-iMG proximity in iMG-_diss_RO. **h**, Surface rendering of iMG for CTRL (top) and poly(I:C) (bottom). BRN3^+^-spots color-coded based on the proximity to the iMG surface with spots < 5 µm (magenta) and spots > 5 µm (cyan). Scale bar: 10 µm. **i**, Boxplot of percent of magenta BRN3^+^-spots. Symbols: single ROI of three biological replicates from five independent differentiations. Wilcoxon rank-sum test. For detailed statistical analysis, see **Supplementary Table 4**. ***p < 0.001. * p < 0.05. ^ns^p > 0.05, not significant. Abbreviations: BRN3: brain-specific homeobox/POU domain protein 3B. CTRL: untreated control. iMG: microglia-like cells. iMG-_diss_RO: microglia integrated into dissociated retinal organoid culture. Poly(I:C): polyinosinic: polycytidylic acid. preMG: microglia precursor cells. _diss_RO: dissociated retinal organoid cultures. RT-qPCR: real-time quantitative polymerase chain reaction. ROI: region of interest. SEM: standard error of the mean. TLR3: toll-like receptor 3. WK: week after the start of RO differentiation.

### iMG presence influences poly(I:C)-mediated inflammatory secretome signature and cell proliferation

To obtain insights into how iMG presence affects the poly(I:C)-mediated neuro-immune response, we analyzed the supernatant of _diss_RO and iMG-_diss_RO after 24h of poly(I:C) stimulation and compared it to the untreated control (**Figure 4a**).

**Figure 4-.**
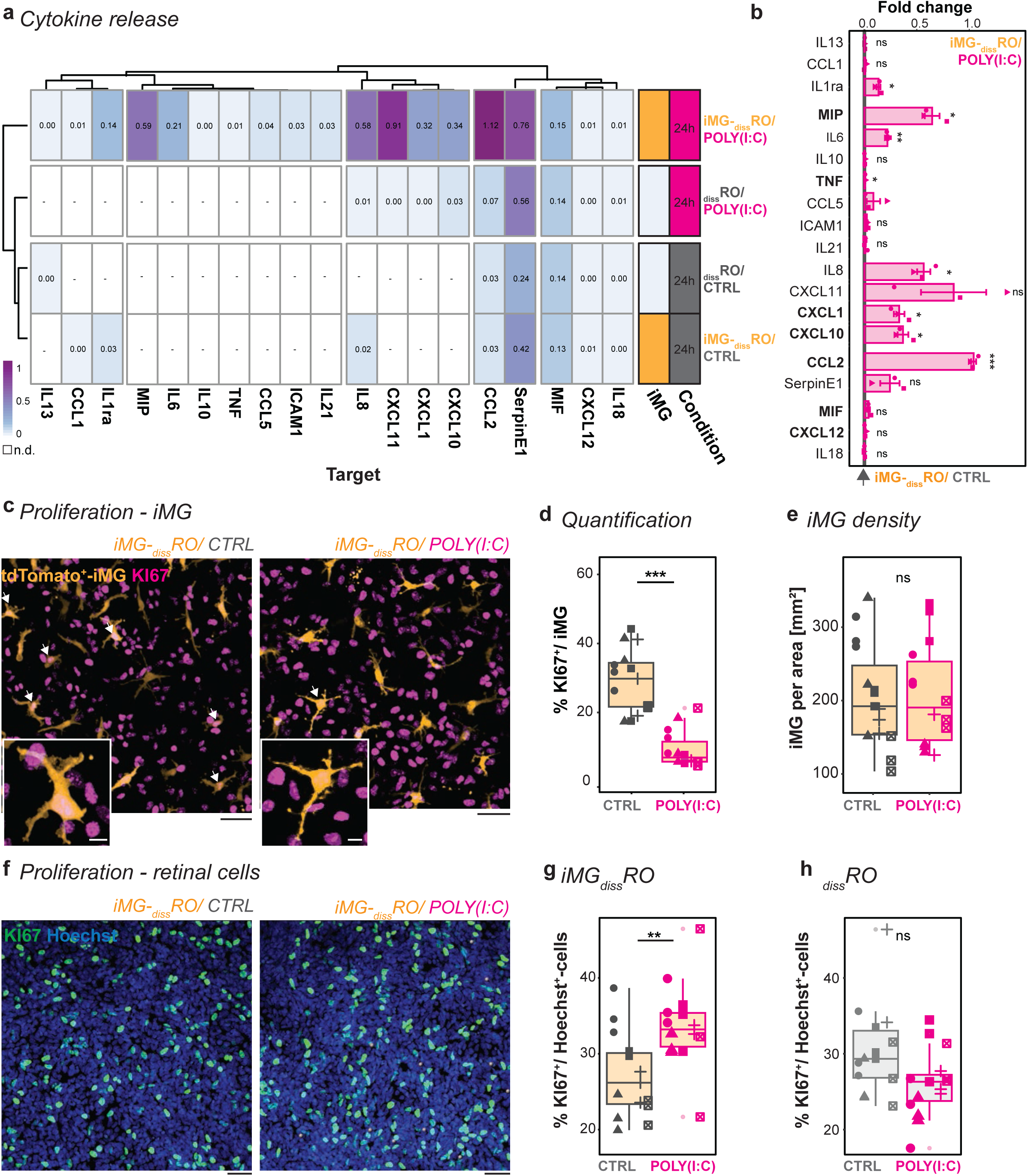
Poly(I:C)-mediated microglia-dependent consequences on the retinal environment. **a-b**, Release of inflammatory cytokines and chemokines into the supernatant based on the experimental paradigm described in Figure 3a for control (CTRL, grey) and poly(I:C) (magenta) after 24-hour stimulation. **a**, Heatmap with color-coded mean pixel intensity relative to the reference of three independent differentiations (for individual plots, see **Supplementary Figure 5a**). White: n.d. Side-bar: condition with iMG (orange) or without (white) or CTRL *versus* poly(I:C). **b**, Bar chart with SEM: Fold change of pixel intensity upon poly(I:C) stimulation relative to CTRL. Each dot is an independent differentiation (n=3). Shapiro-Wilk normality test <0.05, Wilcoxon rank-sum test. Shapiro-Wilk normality test > 0.05, one sample t-test. **c-e**, iMG proliferation rate in iMG-_diss_RO for CTRL (left) and poly(I:C) (right). **c**, Example ROI image of tdTomato^+^-iMG (orange) and immunostained for the proliferation marker KI67 (magenta). White arrow: KI67^+^-expressing iMG. Scale bar: 50µm. Zoom-in: Scale bar: 10 µm. **d**, Boxplot of KI67^+^/iMG percentage. Wilcoxon rank-sum test. **e**, Boxplot of iMG per area. Students’ t-test. **d-e**, Symbols: single ROI of three biological replicates from five independent differentiations. **f-h**, Proliferation of retinal cells excluding iMG in iMG-_diss_RO (**f-g**) and _diss_RO (**h**). **f**, Example ROI images counterstained for the nuclei-dye Hoechst (blue) and immunostained for the proliferation marker KI67 (green) for CTRL (left) and poly(I:C) stimulation (right). Scale bar: 50µm. **g-h**, Boxplot percent of KI67^+^-cells relative to Hoechst^+^-cells for CTRL and poly(I:C) in iMG-_diss_RO (**g**, excluding KI67^+^/iMG) and _diss_RO (**h**, lacking iMG). Symbols: single ROI of three biological replicates from five independent differentiations. Students’ t-test. For detailed statistical analysis, see **Supplementary Table 4**. ***p < 0.001. **p < 0.01. *p < 0.05. ^ns^p > 0.05, not significant. Abbreviations: CTRL: untreated control. iMG: microglia-like cells. iMG-_diss_RO: microglia integrated into dissociated retinal organoid culture. KI67: marker of proliferation KI-67. n.d.: not detectable. Poly(I:C): polyinosinic: polycytidylic acid. _diss_RO: dissociated retinal organoid cultures without iMG. ROI: region of interest. SEM: standard error of the mean. WK: week after the start of RO differentiation.

At baseline without poly(I:C) stimulation, _diss_RO with or without iMG were comparable, showing a similar set of secreted mediators, including MIF, CCL2, CXCL12, IL18, and SerpinE1 (**Figure 4a**, **Supplementary Figure 5a-b**). _diss_RO exposed to poly(I:C) formed a separate cluster with only moderate differences from the controls. The additional detected cytokines CXCL10, CXCL11, CXCL1, and IL8 belong to the CXC family and are known to be secreted by astrocytes [91–93]. Indeed, we verified the presence of GFAP^+^-glia cells in _diss_RO (**Supplementary Figure 5c**). The most robust inflammatory secretome signature occurred when we stimulated iMG-_diss_RO with poly(I:C). The previous four factors were significantly higher released, and we detected an additional eight secreted inflammatory mediators such as TNFα, IL6, and MIP (**Figure 4a-b**). Since those factors have already been partially upregulated in _diss_RO, iMG seemed to amplify the signal. On a note, approximately half of the inflammatory mediators assayed were not secreted in any condition (**Supplementary Figure 5d**), and also not after only 2- or 4 hours of poly(I:C) exposure in iMG-_diss_RO (**Supplementary Figure 5e**).

CCL2 (C-C Motif Chemokine Ligand 2) has been one of the strongest upregulated factors upon poly(I:C) stimulation in iMG-_diss_RO (**Figure 4b**). Besides being involved in the homing of monocytes and T-cells from the periphery [94, 95], CCL2 also contributes to neuronal proliferation in concert with the other upregulated secreted factors CXCL12, MIF, MIP, TNFα, CXCL1, and CXCL10 [28, 30, 96–99] (**Figure 4b**). To investigate the consequences on the number of proliferating cells, we immunostained for the proliferation marker KI67 (**Figure 4c-h**). In iMG-_diss_RO, poly(I:C) stimulation significantly reduced the number of KI67^+^/iMG-cells compared to the control (**Figure 4c-d**), which supports data in rodents [100]. Yet, the overall iMG density remained similar (**Figure 4e**), suggesting that iMG are less proliferative. In contrast, the overall number of proliferating retinal cells excluding iMG significantly increased upon poly(I:C) stimulation in both iMG-_diss_RO (**Figure 4f-g**) and iMG-RO (**Supplementary Figure 6a**). This effect aligns with rodent studies after prenatal immune challenges [16, 22, 101]. Since the secretion of proliferation-associated factors was only upregulated in iMG-_diss_RO, we determined the number of KI67^+^/Hoechst^+^-cell in _diss_RO without iMG. Due to the lack of increased proliferation-associated factors and iMG presence, poly(I:C) failed to increase KI67^+^-cells (**Figure 4h**), emphasizing an iMG-dependent effect on cell proliferation upon poly(I:C) exposure.

Next, to identify whether CCL2 is the primary mediator of this effect, we applied 10 ng/ml CCL2 to iMG-_diss_RO cultures at WK20 and analyzed the consequences 24h later (**Supplementary Figure 6b**). In contrast to poly(I:C) stimulation, CCL2 exposure did not increase the overall proliferation rate (**Supplementary Figure 6c**). Even if we applied higher CCL2 concentrations, the ratio of KI67^+^/Hoechst^+^-cells remained the same. Unexpectedly, the ratio of KI67^+^/iMG rose with 10 ng/ml CCL2 (**Supplementary Figure 6d**), which is in contrast to the poly(I:C) condition (**Figure 4c-d**). This suggests that CCL2 alone cannot drive the observed phenotypes and that the interplay with additional proliferation-associated factors is critical.

### Ibuprofen dampens poly(I:C)-induced iMG phenotypes and reduces cell proliferation

Besides cytokines and chemokines, another hallmark of inflammation is the secretion of prostaglandins such as PGE2, which mediate classic symptoms of inflammation [102, 103]. Indeed, we found that iMG-_diss_RO stimulated with poly(I:C) for 24h showed increased PGE2 levels in the supernatant (**Figure 5a-b**). NSAIDs like ibuprofen target the enzymes cyclooxygenase 1 and 2 (*PTGS1*/COX1, *PTGS2*/COX2, respectively) and prevent arachidonic acid conversion into prostaglandins [70, 71]. Simultaneous exposure of poly(I:C) with the active enantiomer S(+)-ibuprofen dampened PGE2 upregulation (**Figure 5b**). Next, we investigate the release of cytokine and chemokine into the supernatant during the poly(I:C)-mediated neuro-immune challenge when we simultaneously applied S(+)- ibuprofen. Most inflammatory mediators remained unaffected upon exposure to S(+)- ibuprofen (**Supplementary Figure 7a**). Only TNF secretion increased (**Figure 5c**), which aligns with a previous study identifying that PGE2 inhibits TNF expression in macrophage cell lines *in vitro* [104].

**Figure 5-.**
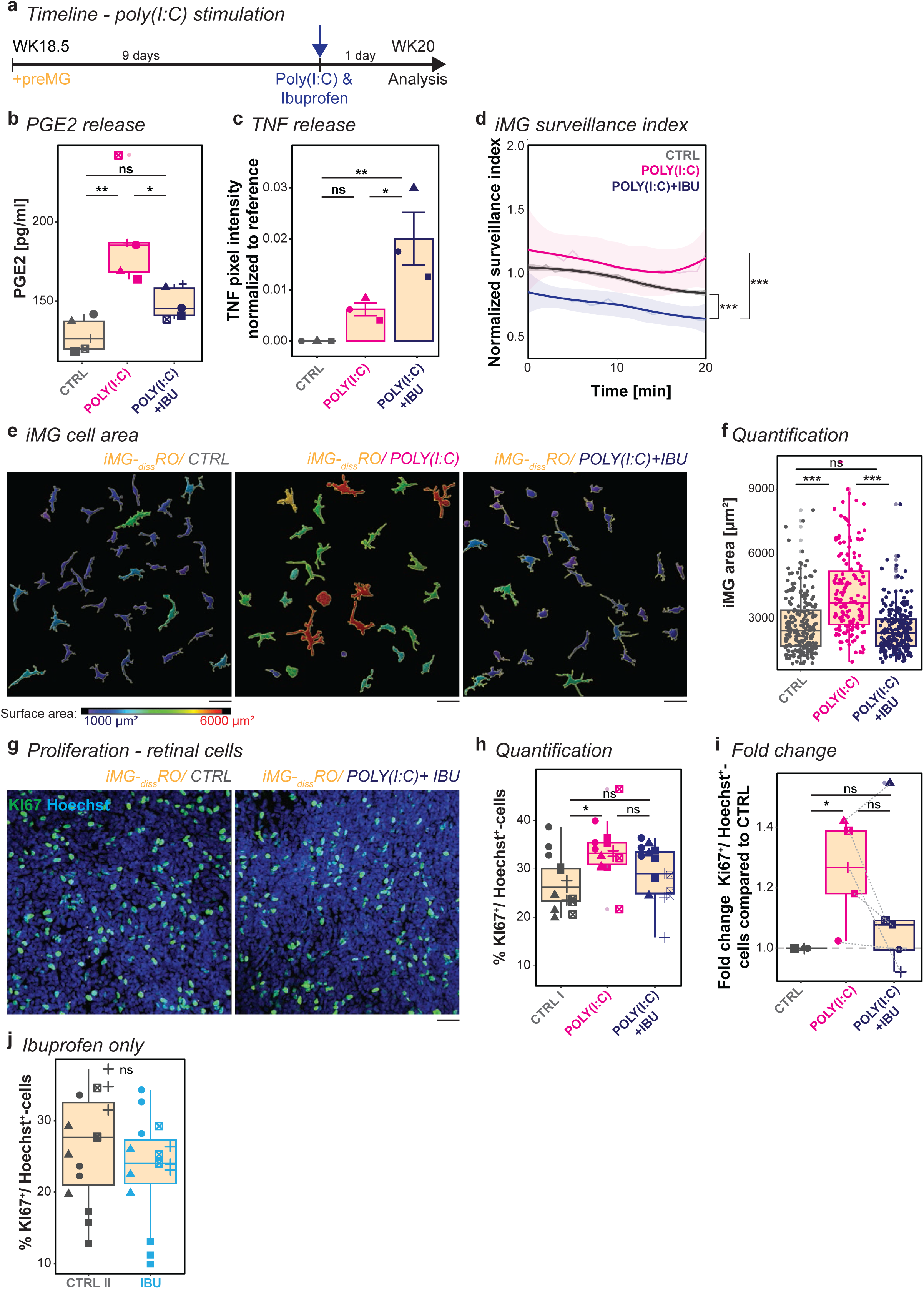
Ibuprofen partially reverses poly(I:C)-mediated consequences on iMG and cell proliferation. **a**, Experimental timeline. At WK18.5, preMG are added to _diss_RO. After nine days, cultures received fresh medium for control (CTRL, grey), poly(I:C) (magenta), or poly(I:C) and S(+)- ibuprofen (poly(I:C)+IBU, blue) for 24 hours before analysis. **b**, PGE2 is released into the supernatant of iMG-_diss_RO. Boxplot of PGE2 concentration [pg/ml] after CTRL, poly(I:C), and poly(I:C)+IBU. Each symbol: an independent differentiation (n=5). One-way ANOVA with post-hoc Tukey’s test. **c**, Release of TNF into the supernatant of iMG-_diss_RO. Boxplot of pixel intensity normalized to reference for CTRL, poly(I:C), and poly(I:C)+IBU. Each symbol is an independent differentiation (n=3). One-way ANOVA with post-hoc Tukey’s test. **d**, iMG-_diss_RO live imaging for 20 minutes after 24h stimulation. iMG surveillance index normalized to the mean surveillance of the cells in CTRL with a 95% confidence interval. Data from five independent differentiations. Kruskal-Wallis test with post-hoc Dunn’s test. **e-f**, iMG surface area in iMG-_diss_RO. **e**, iMG surface rendering for CTRL (left), poly(I:C) (middle), and poly(I:C)+IBU (right), color-coded based on surface area: blue = 1000 µm^2^ to red = 6000 µm^2^. Scale bar: 50 µm. **f**, Boxplot of individual iMG surface areas. iMG from five independent differentiations. Kruskal-Wallis test with post-hoc Dunn’s test. **g-i**, Proliferation rate of retinal cells in iMG-_diss_RO excluding iMG. **g**, Example ROI of counterstained nuclei-dye Hoechst (blue) and immunostained KI67 (green) for CTRL (left) and poly(I:C)+IBU (right). Scale bar: 50 µm. **h**, Boxplot percent of KI67^+^-cells relative to Hoechst^+^-cells for CTRL, poly(I:C), and poly(I:C)+IBU. Symbols: single ROI of three biological replicates from five independent differentiations. One-way ANOVA with post-hoc Tukey’s test. **i**, Fold change of median percent of KI67^+^-cells relative to Hoechst^+^-cells compared to CTRL. For each condition, three ROIs per biological replicate. One sample t-test and Student’s t-est. **j**, Boxplot percent of KI67^+^-cells relative to iMG in iMG-_diss_RO for CTRL and only IBU exposure (light-blue) for 24 hours. Symbols: single ROI of three biological replicates from five independent differentiations. Students’s t-test. For detailed statistical analysis, see **Supplementary Table 4**. ***p < 0.001. **p < 0.01. *p < 0.05. ^ns^p > 0.05, not significant. Abbreviations: CTRL: untreated control. _diss_RO: dissociated retinal organoid cultures without iMG. IBU: S(+)-ibuprofen. iMG: microglia-like cells. iMG-_diss_RO: microglia integrated into dissociated retinal organoid culture. KI67: marker of proliferation KI-67. PGE2: prostaglandin E2. Poly(I:C): polyinosinic: polycytidylic acid. Poly(I:C)+IBU: poly(I:C) and S(+)-ibuprofen. preMG: microglia precursor cells. ROI: region of interest. TNF: tumor necrosis factor alpha. WK: week after the start of RO differentiation.

Since microglia have been shown to constitutively express PTGS1/COX1 [105]and ibuprofen targets PTGS1/COX1, we revisited iMG surveillance and monitored their activity (**Supplementary Video 3**). 24h following exposure of S(+)-ibuprofen simultaneously with poly(I:C), iMG surveillance significantly reduced compared to just poly(I:C) and even below the control level in _diss_RO (**Figure 5d**). Morphologically, iMG remained confined, exhibiting less cell surface area than just poly(I:C) exposure (**Figure 5e-f**), indicating a dampened activity and underlining a direct ibuprofen-mediated effect on iMG.

To further examine whether ibuprofen improves the consequences of the prenatal neuro-immune challenge, we revisited the increased proliferation phenotype upon poly(I:C) exposure. Following simultaneous treatment with S(+)-ibuprofen, the ratio of KI67^+^/Hoechst^+^-cells reduced for four out of five differentiation in iMG-_diss_RO compared to poly(I:C) stimulation alone (**Figure 5g-i**). Since ibuprofen has been associated with anti-proliferative effects in cancer cell lines [106, 107], we evaluated S(+)-ibuprofen without poly(I:C) exposure. The number of proliferating cells remained similar (**Figure 5j**). In iMG-RO, we observed a similar beneficial effect, emphasizing the comparability of the 2D and the 3D (**Supplementary Figure 7b-c**).

### Ibuprofen depends on iMG to reverse poly(I:C)-mediated effects on neuronal activity

Initially, we described that the OPL formation aligned with iMG integration. Once the OPL is formed, spontaneous glutamatergic activity shapes neuronal circuits *in vivo* [108]. Since the iMG-_diss_RO expressed synaptic markers (**Figure 6a**), we visualized spontaneous calcium transients as a correlate for neuronal activity. First, we transduced _diss_RO with adeno-associated virus (AAV), which is independent of TLR3-signaling [109]. The AAV encoded for the calcium sensor GCAMP6s under the control of the ubiquitous EF1α promoter [10, 110], resulting in a broad expression across retinal cell types (**Supplementary Figure 8a-e**). Importantly, to exclude any AAV-mediated microglia activation, we applied the virus to _diss_RO at WK17 and 1.5 weeks before adding preMG (**Figure 6b**). At WK20, we analyzed the spontaneous calcium transients (**Figure 6c**, **Supplementary Video 4**). The calcium peak amplitude and the mean frequency remained similar in _diss_RO and iMG-_diss_RO (**Figure 6d-e**). The calcium transients were either abolished after application of the voltage-gated sodium channel blocker tetrodotoxin (TTX) (**Figure 6f**, **Supplementary Video 5**) or significantly reduced after pharmacological blocking of glutamatergic synaptic transmission using a combination of CPP, NBQX, and APB (**Figure 6g**, **Supplementary Video 6**). Furthermore, the spontaneous calcium activity depended on extracellular calcium because the transients stopped when we applied the Ca^2+^-chelator EGTA into the media (**Figure 6h**, **Supplementary Video 7**).

**Figure 6-.**
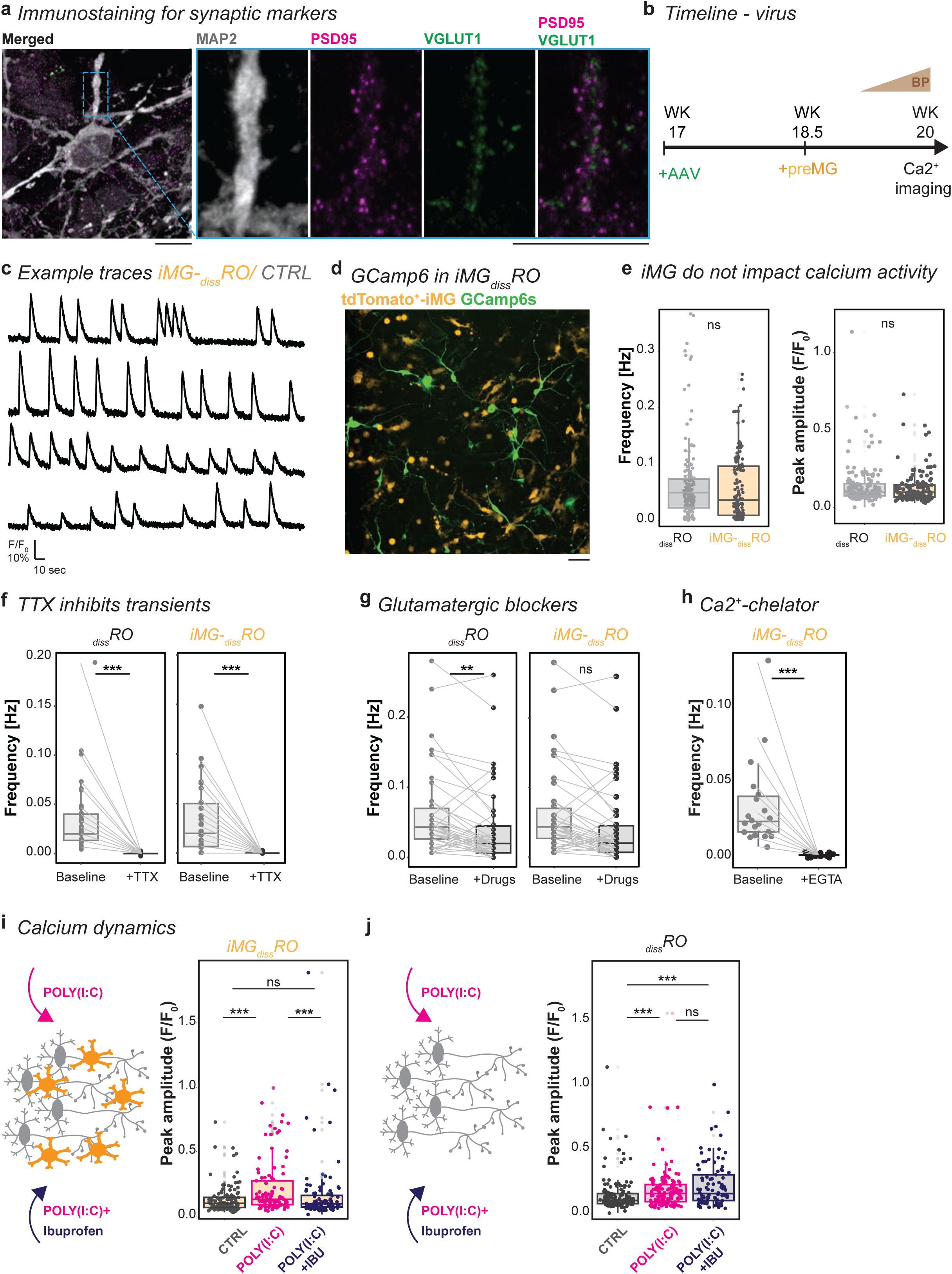
Calcium dynamics are affected upon poly(I:C), yet ibuprofen needs iMG to reverse the phenotype. **a**, Example image of iMG-_diss_RO immunostained for the neuronal marker MAP2 (grey), the presynaptic marker VGLUT1 (green), and the postsynaptic marker PSD95 (magenta) with zoom-in. Scale bar: 10 µm. **b**, Experimental timeline. At WK17, _diss_RO transduced with AAV2-GCAMP6s. preMG added at WK18.5. Four days before calcium imaging, gradual transition to Brain-Phys medium until WK20. **c**, Example traces of spontaneous calcium transients in iMG-_diss_RO. **d**, Example ROI image of iMG-_diss_RO expression of the calcium sensor GCAMP6s (green) and tdTomato^+^-iMG (orange). Scale bar: 50µm. **e**, Spontaneous calcium dynamics during five minutes recording in _diss_RO (grey) and iMG-_diss_RO (orange). Boxplot of the mean frequency [Hz] (left) and the mean peak amplitude [F/F_0_] (right). Wilcoxon rank-sum test. **f**, TTX abolishes calcium transients in _diss_RO (left) and iMG-_diss_RO (right). Boxplot of mean frequency [Hz] during 150-sec baseline recording (baseline) following TTX application for another 150 sec (+TTX). One-sample Wilcoxon signed rank test. **g**, Boxplot of mean frequency [Hz] during 150-sec baseline recording (baseline) following exposure to glutamatergic blockers CPP, APB, and NBQX for 150 sec in _diss_RO (left) and iMG-_diss_RO (right). Wilcoxon rank-sum test. **h**, Boxplot of mean frequency [Hz] during 150-sec baseline recording (baseline) following Ca2^+^-chelator EGTA application for another 150 sec in iMG-_diss_RO. One-sample Wilcoxon signed rank test. **i-j**, Spontaneous calcium dynamics during five minutes recording after 24 hours of either fresh medium for control (CTRL, grey), poly(I:C) (magenta), or poly(I:C) and S(+)-ibuprofen (poly(I:C)+IBU, blue) exposure in iMG-_diss_RO (**i**) and _diss_RO (**j**). Boxplot of the mean peak amplitude [F/F_0_]. Each dot represents an active cell. Recordings from five independent differentiations. Kruskal-Wallis test with post-hoc Dunn’s test. For detailed statistical analysis, see **Supplementary Table 4**. ***p < 0.001. **p < 0.01. ^ns^p > 0.05, not significant. Abbreviations: AAV: adeno-associated virus. APB: 2-aminoethoxydiphenyl borate. CPP: 4-(3-phosphonopropyl)piperazine-2-carboxylic acid. CTRL: untreated control. _diss_RO: dissociated retinal organoid cultures without iMG. EGTA: Ethylene glycol tetraacetic acid. IBU: ibuprofen. iMG: microglia-like cells. iMG-_diss_RO: microglia integrated into dissociated retinal organoid culture. MAP2: microtubule-associated protein 2. NBQX: 2,3-dioxo-6-nitro-7-sulfamoyl-benzo[f]quinoxaline. Poly(I:C): polyinosinic: polycytidylic acid. PSD95: postsynaptic density protein 95. _diss_RO: dissociated retinal organoid cultures. Sec: seconds. TTX: tetrodotoxin. VGLUT1: vesicular glutamate transporter 1. WK: week after the start of RO differentiation.

Inflammatory factors have been shown to modulate neuronal activity [111–114]. Indeed, we found that poly(I:C) exposure in both _diss_RO and iMG-_diss_RO significantly increased the calcium peak amplitude of individual cells (**Figure 6i-j**) and had no effect on the mean frequency (**Supplementary Figure 8f**). However, when we simultaneously applied S(+)-ibuprofen, strikingly, the peak amplitude was only restored to the control condition in iMG-_diss_RO (**Figure 6i**). In _diss_RO, the peak amplitude remained elevated compared to the control (**Figure 6j**), and the frequency remained the same (**Supplementary Figure 8f**). This data suggests that iMG presence is critical for the effect of ibuprofen on restoring poly(I:C)-induced changes in the calcium dynamics.

### Both ibuprofen targets, PTGS1 in microglia and PTGS2, are required to rescue calcium dynamics

To identify the mechanism behind the microglia-dependent response upon ibuprofen exposure, we revisited the two main targets of ibuprofen, *PTGS1* and *PTGS2*, at the transcriptional level. In the RO sequencing data [10], *PTGS2* transcripts occurred in Müller glial and horizontal cells, whereas *PTGS1* transcripts were absent (**Supplementary Figure 9a**). Since this dataset lacks iMG, we collected RO with and without iMG around WK20 and analyzed the mRNA transcript levels of both enzymes. *PTGS1* relative to *GAPDH* was significantly increased in iMG-RO compared to RO (**Figure 7a**), whereas the *PTGS2* levels remained similar (**Figure 7b**).

**Figure 7-.**
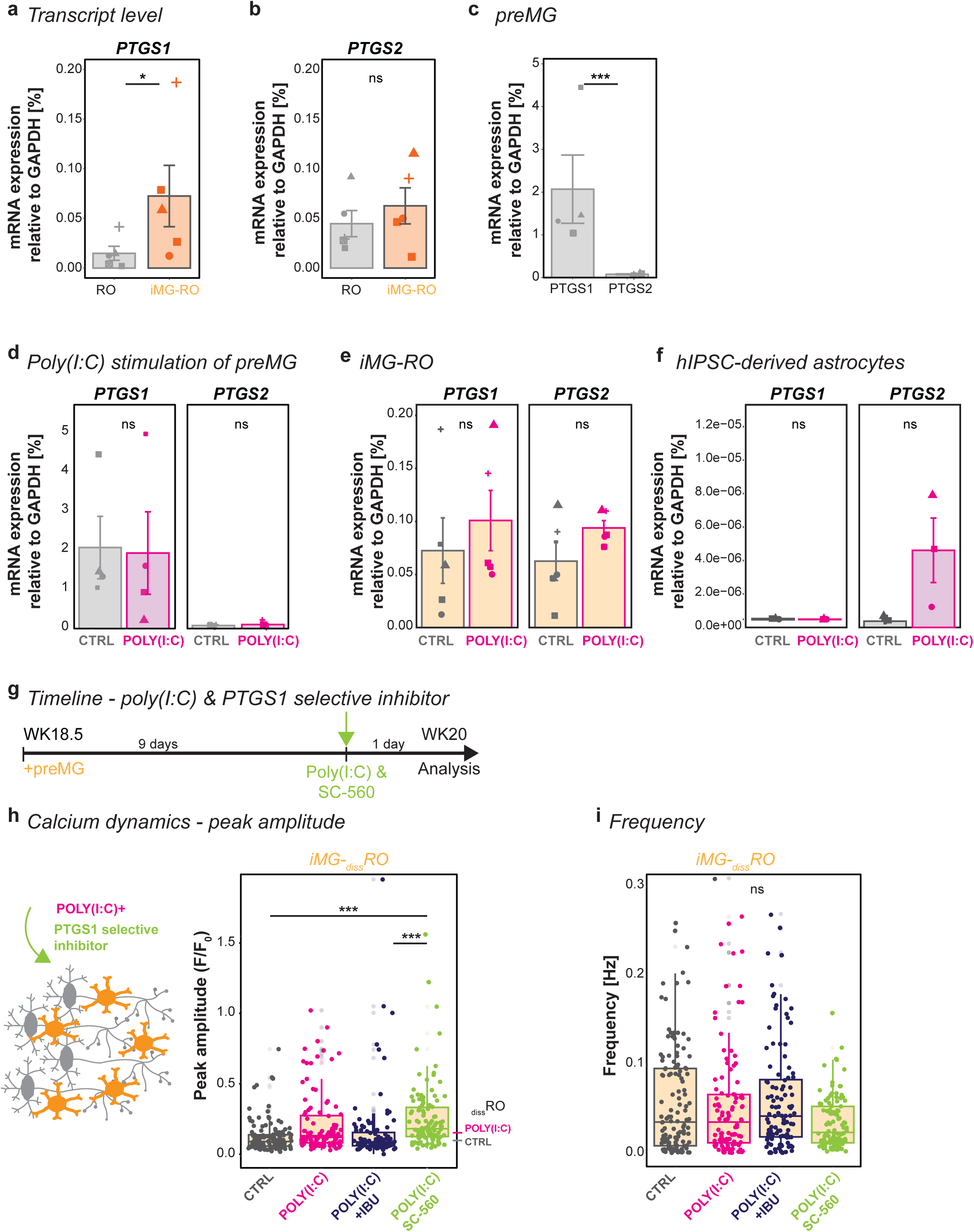
Cell type-dependent transcriptional differences in PTGS1 and PTGS2 point to iMG impact on calcium dynamics. **a-f**, RT-qPCR. Bar chart with SEM of mean mRNA transcript expression relative to GAPDH for PTGS1 (**a**, **c-f**) and PTGS2 (**b-f**) for RO *versus* iMG-RO (**a-b**), for preMG (**c-d**), for iMG-RO (**e**), and hIPSC-derived astrocytes (**f**) for untreated (**a-c**) and CTRL *versus* poly(I:C) (**d-f**). **g**, Experimental timeline similar to Figure 6b with poly(I:C) and SC-560 exposure for 24h in iMG-_diss_RO at WK20. **h-i**, Spontaneous calcium dynamics in iMG-_diss_RO during 5 minutes recording after 24 hours of either fresh medium for control (CTRL, grey), poly(I:C) (magenta), poly(I:C) and S(+)-ibuprofen (poly(I:C)+IBU, blue), or poly(I:C) and SC-560 (poly(I:C)+SC-560, green) exposure. Boxplot of mean peak amplitude [F/F_0_, **h**] and mean frequency [Hz, **i**]. Each dot represents an active cell from five independent differentiations. Peak amplitude: reference lines for the median of the control and poly(I:C) stimulation in _diss_RO from Figure 6j. Kruskal-Wallis test with post-hoc Dunn’s test. For detailed statistical analysis, see **Supplementary Table 4**. ***p < 0.001. *p < 0.05. ^ns^p > 0.05, not significant. Abbreviations: CTRL: untreated control. GAPDH: glyceraldehyde 3-phosphate dehydrogenase. H: hour. hIPSC: human induced pluripotent stem cell. IBU: ibuprofen. iMG: microglia-like cells. iMG-_diss_RO: microglia integrated into dissociated retinal organoid culture. Poly(I:C): polyinosinic: polycytidylic acid. preMG: microglia precursor cell. PTGS1: prostaglandin-endoperoxide synthase 1 (COX1), PTGS2: prostaglandin-endoperoxide synthase 2. RT-qPCR: real-time quantitative polymerase chain reaction. SC-560: PTGS1/COX-1 selective inhibitor. SEM: standard error of the mean. WK: week after the start of RO differentiation.

Microglia have been shown to express PTGS1 constitutively [105], and a common assumption is that PTGS2 is upregulated during inflammatory conditions [115]. To validate these assumptions, we investigated the transcript level in preMG cultures. Indeed, the absolute expression of *PTGS1* compared to *GAPDH* was significantly higher than that of *PTGS2* (**Figure 7c**). Yet, neither PTGS1 nor PTGS2 levels increased upon poly(I:C) exposure following poly(I:C) stimulation (**Figure 7d**). At the same time, we observed a trend for *PTGS1* and *PTGS2* increase in iMG-RO (**Figure 7e**). When we analyzed the cytokine assay (**Figure 4a-b**), we observed several secreted factors known to be released by astrocytes. In RO, astrocytes/Mueller-glia show a low expression level of TLR3 (**Supplementary Figure 9b**). We confirmed this expression in hIPSC-derived astrocyte cultures [116], which further upregulate *TLR3* transcripts upon poly(I:C) exposure (**Supplementary Figure 9c**). When we profiled *PTGS1* and *PTGS2*, we found that hIPSC-derived astrocytes significantly increased *PTGS2* transcripts (**Figure 7f**). This emphasizes that preMG and astrocytes differentially express the enzymes cyclooxygenase 1 and 2, respectively.

Since S(+)-ibuprofen targets both enzymes simultaneously under poly(I:C) exposure, their interaction balance is not disturbed in contrast to _diss_RO, where iMG are missing. To test this hypothesis, we took advantage of the inhibitor SC-560, which has a 700-fold selectivity for PTGS1 over PTGS2 [117, 118] and, therefore, will directly affect iMG in iMG-_diss_RO. When we applied SC-560 together with poly(I:C) for 24 hours (**Figure 7g**) and measured the calcium dynamics in iMG-_diss_RO, the calcium peak amplitude was similarly upregulated as in _diss_RO following ibuprofen treatment (**Figure 7h**). The frequency remained unaffected (**Figure 7i**). This suggests that inhibition of functional PTGS1 or the simple lack of iMG prevents the positive effect of ibuprofen from restoring the calcium peak amplitude, highlighting a critical interplay between PTGS1-expressing iMG and PTGS2-mediated poly(I:C) upregulation in other cells.

## Discussion

This study highlights the importance of including microglia in hIPSC-neuronal organoids to mimic fetal brain development and adequately model pathogen- and ibuprofen-mediated processes.

### Prenatal neuro-immune challenge

The immune system protects a pregnant woman and her offspring from diseases [119]. Yet, if the mother is exposed to a TORCH infection (Toxoplasmosis, Others, Rubella, Cytomegalovirus, Herpes), this can cause a threat to the fetus [2, 3]. Through vertical transmission, TORCH can directly harm the developing fetus, inducing growth restrictions or birth defects such as blindness [120]. For example, the Zika virus has been shown to target human fetal microglia [121] and to activate the innate immune receptor TLR3 [46, 122]. Thus, we mimicked the TLR3 signaling pathway using poly(I:C) [65]. *In vivo*, the receptor is expressed predominantly in microglia but also in astrocytes, endothelial cells, and pericytes, while only minimally in neurons and neuronal progenitors [51]. We confirmed this expression pattern in preMG (**Figure 3b**) and hIPSC-derived astrocyte cultures (**Supplementary Figure 9c**). Furthermore, we demonstrated that 24 hours of poly(I:C) exposure resulted in a robust microglia-dependent cytokine release (**Figure 4a-b**), accompanied by enhanced iMG surveillance (**Figure 3c**), elevated calcium peak amplitude in retinal cells (**Figure 6i**), and increased retinal cell proliferation (**Figure 4f-g**). Our observations in _diss_RO lacking iMG showed a trend of reduced proliferation (**Figure 4h**). This aligns with rodent models in which the proliferation rate remained unaffected if microglia were depleted and the pregnant females were exposed to an immune challenge during embryonic and early postnatal stages [22]. Contrarily, the proliferation increased in the presence of microglia [16, 22, 101], comparable to our data (**Figure 4f-g**). Microglia-dependent effects have also been reported on the transcriptional level after prenatal neuro-immune challenges, such as IFNɣ stimulation of brain organoids [123] or three days of poly(I:C) exposure on E15.5 rodent brain [51], emphasizing the microglia relevance in the inflammatory response.

This raises the question of how to treat infectious diseases during pregnancy. Our insights are limited to how a viral infection affects the fetus and whether treatments are appropriate to prevent adverse pregnancy outcomes [119]. NSAIDs act on prostaglandin G/H synthase 1 and 2 (PTGS1/COX1 and PTGS2/ COX2), resulting in anti-inflammatory, antipyretic, and analgesic properties [124, 125]. We decided on ibuprofen in this study because Germany allows this medication during the first two trimesters of pregnancy [69]. Ibuprofen crosses the placental barrier and accesses the developing fetus [126, 127]. We found that by applying ibuprofen simultaneously with poly(I:C), we could ameliorate the increased cell proliferation (**Figure 5g-i**), restore the calcium peak amplitude (**Figure 6i**), and dampen the poly(I:C)-mediated microglial response (**Figure 5d-f**). Interestingly, ibuprofen did not reduce most of the secreted inflammatory mediators in iMG-_diss_RO (**Supplementary Figure 7a**). This could be due to our culture model, which lacks infiltrating immune cells to resolve the inflammatory response.

Ibuprofen might directly interfere with microglia, which constitutively express PTGS1 over PTGS2 (**Figure 7c**) and do not show a difference in the expression level upon poly(I:C) stimulation (**Figure 7d**). When we selectively inhibited PTGS1 with SC-560, the calcium dynamics did not recover (**Figure 7h**), similar to _diss_RO. These results emphasize the importance of including all relevant cell types, as PTGS1 and PTGS2 interaction between microglia and retinal cell types is crucial for the beneficial effects of ibuprofen.

### The effects of microglia in retinal organoids

Recently, three studies have been published focusing on iMG integration into retinal organoids [39–41]. Both Usui-Ouchi *et al.* and Chichagova *et al.* performed their integration and investigation on a timeline similar to our study and compared RO with and without iMG. Usui-Ouchi *et al.* confirmed the iMG integration into the developing OPL (**Figure 1e-f**).

Interestingly, they show that the integration temporarily increases the pro-inflammatory factors *IL1B*, *TNF,* and *IL6* using qRT-PCR. Our cytokine release assay did not detect differences in IL6, TNF, and IL1B between _diss_RO and iMG-_diss_RO (**Figure 4a**, **Supplementary Figure 5a-b**, **d**), and only after poly(I:C) stimulation, IL-6 and TNF were released in iMG-_diss_RO. We also performed qRT-PCR for these three pro-inflammatory markers and observed a similar mRNA upregulation as the authors describe. However, poly(I:C) stimulation exceeded at least 500-fold (data not shown). This indicates that translation from mRNA to an actual release into the supernatant might be tightly controlled. Chichagova *et al.* mimicked bacterial infection with lipopolysaccharide (LPS) for 24 hours and validated inflammatory mediators in the supernatant using a different assay. They found differences in the LPS response when iMG are present in the RO, but interestingly, there is no effect on the CCL2/MCP-1, one of our strongest affected chemokines (**Figure 4a**).

Furthermore, they should have included the control conditions and could miss potential changes due to the iMG integration, as reported before by Usui-Ouchi *et al*. Finally, the study by Gao *et al.* compares mostly macrophages not integrated into a neuronal environment, which we named in our study preMG, after either LPS and poly(I:C) stimulation. We consider this comparison suboptimal due to the known effects of LPS and poly(I:C) on neuronal populations [16, 22, 101], also exemplified in the impact on the peak amplitude of our calcium dynamics (**Figure 6i-j**). The authors suggest that in their iMG co-cultured with D30-old RO, a similar reduction in BRN3^+^-cells as we have seen (**Figure 2g**). Yet, they do not show iMG-BRN3^+^-interaction (**Figure 2h**) and only describe in the text that they found a few microglia in the retina without insights into their positioning, making it challenging to derive further conclusions.

Overall, the above studies are inconclusive regarding the ideal timing for studying microglia-neuron interaction and the consequences of prenatal neuro-inflammation and drug-mediated effects. iMG-RO guided us to WK20, where iMG integration matched human fetal tissue (**Figure 1c**). Yet, we were challenged with the ganglion cell loss (**Figure 2a-b**), which we overcame with our 2D-_diss_RO model that reflected a similar cell type composition as RO (**Figure 2c-d**). Integrated iMG were functional, recapitulating developmental tasks such as regulating ganglion cell number (**Figure 2e**) and removing cellular debris (**Supplementary Figure 4d-e**). Furthermore, this model enabled us to compare different treatment paradigms of the same RO origin. We distributed a pool of dissociated ROs across multiple wells, circumventing common organoid-to-organoid variability. This enabled us to go beyond investigating the role of iMG under ‘healthy’ conditions [39, 41, 128] and to specifically target the effects of iMG following a prenatal neuro-immune challenge and an anti-inflammatory treatment (**Figure 3-6**). Our results highlight the importance of integrating microglia into organoids to study embryonic development.

Organoids commonly lack microglia and consequently dismiss their impact on neurodevelopmental and neurodegenerative diseases [129–133]. In recent years, genetic risk variants have been identified to be enriched or exclusively expressed by microglia [134–136], and inflammation itself has been shown to initiate and drive disease progression [137]. We show that iMG are critical to promoting an inflammatory response, and if the effects of particular prenatal neuro-immune challenges should be tested, this requires iMG. In light of replacing *in vivo* models for drug screening and validation for FDA drug approval [138] with *in vitro* models [139], this becomes critical to achieve drug efficiency in screenings. The limitation of these models to replicate adequately an inflammatory response will severely affect the information regarding the safety of medications during pregnancy for both the pregnant woman and the fetus and will raise serious public health concerns.

In conclusion, our study provides a baseline for the neurodevelopmental role of microglia, the cross-talk with their neuron-glia environment, and how prenatal neuro-immune activation and anti-inflammatory drug treatment are affected. Specifically, our model can serve as a platform for follow-up investigations on drug tests or the interplay between inflammation and microglia activation leading to neurological phenotypes in adulthood. For example, a rubella infection during pregnancy is one of the risk factors for developing schizophrenia [140].

## Limitation

This study focuses on the acute response to an early-life neuro-immune challenge and how microglia contribute to the observed consequences (**Figure 3**). We did not further investigate potential long-term effects caused by, *e.g.*, the increased proliferation rate (**Figure 4f-g**) or the elevated calcium peak amplitude (**Figure 6i**). Moreover, it would be interesting to prolong the stimulation period as infections usually last longer than our tested 24 hours. We focused on WK20 due to the observed microglia-neuron interaction at this single developmental time point (**Figure 1e-g**). As the timing of the immune challenge contributes to differences in the observed outcomes [141, 142], it would be interesting to analyze the effect of TLR3 activation at different developmental time points. Finally, future models must expand on the possibility of including the blood-brain barrier into the system. Identified factors such as MIP, CCL5, CXCL1, and CCL2 (**Figure 4a**) are known candidates for homing monocytes and T-cells from the periphery [95, 143–146], and might be needed to downregulate the inflammatory signature once recruited.

## Material and Methods

### Ethical approval

The ISTA Ethics Officer and Ethics Committee approved the usage of human induced pluripotent stem cells (hIPSC).

### Cell lines

We used two human induced pluripotent stem cell lines (hIPSC): SC 102A-1 GVO-SBI Human Fibroblast-derived (feeder-free) IPSC line (BioCat; hPSCreg.eu: SBLi006-A; in this study referred to SC102A) and the human fibroblast-derived IPSC line 01F49i-N-B7 (Renner lab [10], in this study referred to F49B7). For more details, see (**Supplementary Table 1**).

### Cell culture and human induced pluripotent stem cells

#### Matrigel coating

Matrigel (Corning® Matrigel® hESC-Qualified Matrix, *LDEV-Free, Corning, #354277) was used according to the manufacturer protocol with the following modifications: Matrigel aliquots were dissolved in ice-cold X-Vivo 10 chemically defined, serum-free hematopoietic cell medium (Lonza, #BE04-380Q). Dishes were coated for 1 hour at room temperature.

#### Maintenance of human induced pluripotent stem cells (hIPSCs)

hIPSCs were maintained in mTeSR1 medium (STEMCELL Technologies, #85850) on Matrigel-(Corning, #354277) coated 6-well plates (Corning, #3516) cultured at 37°C and 5% CO_2_ in a humidified incubator (BINDER C150). Before reaching 80% confluency, hIPSCs were passaged as small aggregates every 3-4 days using EDTA dissociation buffer composed of 0.5M EDTA (ethylenediaminetetraacetic acid, K.D. Biomedical, #RGF 3130), 0.9 g (w/v) NaCl (Sigma, #5886) in PBS (phosphate buffered saline, calcium/magnesium-free, Invitrogen, #14190), sterile filtered and stored at 4°C according to [147]. The ISTA Molecular Biology Facility regularly tested hIPSCs for mycoplasma using the Multiplex qPCR assay – 16S DNA according to [148].

#### Freezing and thawing of hIPSCs

For freezing, hIPSCs were washed once with DPBS (Thermo Fisher Scientific, #14190250), incubated in EDTA dissociation buffer for 2.5 minutes, detached as small aggregates using mFReezer (STEMCELL Technologies, #05854), and frozen at −80°C. For long-term storage, hIPSCs aliquots were transferred to liquid nitrogen. For thawing, hIPSCs were removed from liquid nitrogen and quickly thawed in a bead bath at 37°C. hIPSCs were transferred into a falcon tube containing mTesR1 medium. The cells were centrifuged (VWR, Mega Star 3.0R) at 200×g for 3 minutes, then resuspended in mTesR1 medium and transferred into one well of a Matrigel-coated 6-well plate.

#### Generation of tdTomato expressing hIPSC lines

To generate ubiquitous tdTomato expressing hIPSC lines, a reporter construct encoding tdTomato under the constitutive enhancer/β-actin (CAG) promoter (2xCHS4-CAG-tdTomato-SV40-2xCHS4, gift from the Knoblich lab [149]) was inserted into the safe-harbor AAVS1 locus using a CRISPR/CAS9 approach as described previously [150]. For nucleofection, hIPSCs were dissociated into single-cell suspension using Accutase (Merck, #SCR005) treatment for 4 minutes. Cells were collected, centrifuged (VWR, Mega Star 3.0R) at 200×g for 3 minutes, and resuspended in mTeSR1 medium. The Human Stem Cell NucleofectorTM Kit 1 (Lonza, #VPH-5012) was applied using 1 million hIPSCs, 3 µg donor plasmid DNA, and 1 ug CRISPR/CAS9 guideRNA (pXAT2 plasmid, Addgene: #80494). After nucleofection, cells were distributed on six wells of a Matrigel-coated 6-well plate. 5-6 days after the transfection, tdTomato expressing hIPSCs were isolated using fluorescent activated cell sorting (FACS, Sony, SH800SFP). Therefore, hIPSCs were collected using Accutase treatment for 4 minutes, centrifuged at 200g for 3 minutes, and resuspended in mTesR1 medium. Using a 100 µm nozzle, 10.000 hIPSCs were sorted and distributed on three wells of a Matrigel-coated 6-well plate. After 4-5 days, 20 to 30 colonies, which evenly expressed tdTomato, were identified using an EVOS imaging system (Thermo Fisher Scientific), manually picked with a 200 µL tip, and transferred into a Matrigel-coated 96-well plate. The colonies were expanded into 24- (Corning, #3527), 12- (Corning, #3512), and 6-well plates (Corning, #3516). For passaging, refer to “*Maintenance of human induced pluripotent stem cells (hIPSCs)”*.

#### Validation of tdTomato expressing hIPSC lines

Half of the hIPSCs were collected for genotyping when splitting colonies from a 24-well plate to a 12-well plate. DNA was extracted using the DNeasy Blood and Tissue kit (QIAGEN, #69504). All reactions were performed using Q5 Hot Start High Fidelity 2x Master Mix (NEB, #M0494S) with 50-100 ng of template DNA per reaction. PCR was performed using the following primers to identify whether the insertion was heterozygous or homozygous: AAVS1_FWD: 5’-TCGACTTCCCCTCTTCCGATG-3’, AAVS1_WT_REV 5’-CTCAGGTTCTGGGAGAGGGTAG-3’ and AAVS1_Insert_REV 5’-GAGCCTAGGGCCGGGATTCTC-3’as described previously[150]. The size of PCR products was analyzed by gel electrophoresis (wildtype allele 1.4 kbp and target allele 1.2 kbp). Clones with correctly targeted homozygous insertions were expanded.

### Differentiation of retinal organoids, astrocytes, and microglia precursor cells

#### Retinal organoid differentiation

3D-retinal organoids were generated as described with the following modifications [9, 10]: On day 0 of the differentiation, colonies of the hIPSC line F49B7 from two wells of a 6-well plate were cut into evenly sized aggregates using a cell-passaging tool (Thermo Fisher Scientific, #23181-010). After detaching, floating aggregates were transferred with a 1250 µL wide orifice pipette (VWR, #613-0737) into one 9-cm Petri dish (Sarstedt, #82.1473), and cultured in mTeSR1 medium supplemented with 10 µM blebbistatin (Sigma, #B0560-5MG). On day 1, 2, and 3, the medium was gradually exchanged with ¼, ½, and 1, respectively, to NIM (neural induction medium: DMEM/F12 (Gibco, #31331-028), 1×N2-supplement (Gibco, #17502-48), 1% (v/v) NEAA Solution (Sigma, #M7145), 2 µg/mL heparin (Sigma, #H3149-50KU)). From day 4 onwards, 10 mL NIM was changed daily. On day 8, embryoid bodies (EB) were equally distributed onto 8 Matrigel-coated 6-cm dish plates (Corning, #3516) (approximately 20-40 number of EBs/cm^2^) and cultured in 3 mL NIM. From day 16 onwards, NIM was replaced to 3:1-medium consisting of 3 parts DMEM (Thermo Fisher Scientific, #31966047) and one-part F12 medium (Ham’s F-12 Nutrient Mix, Thermo Fisher Scientific, #31765-027) supplemented with 2% (v/v) B27 without vitamin A (Thermo Fisher Scientific, #121587-10), 1% (v/v) NEAA solution (Sigma, #M7145), 1% (v/v) penicillin-streptomycin (Thermo Fisher Scientific, #15140122). Media was changed daily. On day 30, optic-cup structures were detached from the 6-cm plate by checkerboard scraping using a 200 µL pipette tip and transferred into a 9-cm Petri dish containing 10 mL 3:1-medium. The medium was changed twice per week. Between D36 and D42, retinal structures displaying a bright stratified neuroepithelium were manually picked using an EVOS imaging system (Thermo Fisher Scientific). 3D-retinal organoids were not dissected to remove non-retinal tissue. From day 42 onwards, 3:1-medium was supplemented with 10% (v/v) heat-inactivated FBS (Thermo Fisher Scientific, #10270-106) and 100 µM taurine (Sigma, #T0625-25G). The medium was changed twice per week. From week 10, 1 µM retinoic acid was added daily (Sigma, #R2625) while the medium was changed twice per week. From week 14, 3D-retinal organoids were cultured in N2-medium consisting of 3 parts DMEM (Thermo Fisher Scientific, #31966047) and one-part F12 medium (Ham’s F-12 Nutrient Mix, Thermo Fisher Scientific, #31765-027) supplemented with 1×N2 supplement (Gibco, #17502-48), 1% (v/v) NEAA solution (Sigma, #M7145), 1% (v/v) penicillin-streptomycin (Thermo Fisher Scientific, #15140122), 10% (v/v) heat-inactivated FBS (Thermo Fisher Scientific, #10270-106), and 100 µM taurine (Sigma, #T0625-25G). The retinoic acid concentration was reduced to 0.5 µM and added daily. Organoids were cultured at 37°C and 5% CO_2_ in a humidified incubator (BINDER C150).

#### Microglia precursor cell differentiation

The differentiation protocol is identical to the “*Retinal organoid differentiation*” section with the following modifications: 2 wells of the hIPSC line tdTomato-SC102A were used to start the differentiation. On day 1, a final concentration of 12.5 ng/mL recombinant human BMP4 (Bone Morphogenetic Protein 4, Peprotech, #120-05) was added to the medium. From day 8 onwards, NIM was changed twice per week. From day 16 until the termination of the differentiation, cultures were maintained in 3:1-medium, with the medium changed twice per week.

#### Harvesting microglia precursor cells (preMG)

From day 36 onwards, preMG were harvested from the supernatant. For this, the supernatant was passed through a 100 µm cell strainer (Corning, ##352360) and collected in a falcon. After centrifugation (VWR, Mega Star 3.0R) at 200g for 3 minutes, the medium was aspirated, and preMG were resuspended in 3:1-medium supplemented with 50 ng/mL recombinant human MCSF (Macrophage Colony Stimulating Factor, BioLegend, #574804). Cells were counted using an automated cell counter (Bio-Rad, #1450102).

#### Neural Progenitor cell (NPC) differentiation

According to the manufacturer’s instructions, neuronal progenitor cells (NPCs) were generated using the STEMdiff^TM^ SMADi Neural Induction Kit (STEMCELL Technologies, #08581). NPCs were expanded in STEMdiff^TM^ Neural Progenitor Medium (STEMCELL Technologies, #05833) and frozen in STEMdiff^TM^ Neural Progenitor Freezing Medium (STEMCELL Technologies, #05838). NPCs were passaged at 1.25×10^5^ cells/cm^2^ weekly with Accutase^TM^ (STEMCELL Technologies, #07920).

#### Astrocyte differentiation

Astrocytes were differentiated as described previously [151] with minor modifications. For astrocyte differentiation, NPCs were dissociated into single cells using Accutase^TM^ (STEMCELL Technologies, #07920) treatment for 5-10 minutes. Cells were centrifuged at 400×g for 5 minutes (VWR, Mega Star 3.0R), and the medium was aspirated. Cells were resuspended in complete astrocyte-medium composed of astrocyte medium (Sciencell, #1801-b), 2% (v/v) heat-inactivated FBS (Thermo Fisher Scientific, #10270-106), 1% (v/v) astrocyte growth supplement (Sciencell, #1852) and 1% (v/v) penicillin-streptomycin (Thermo Fisher Scientific, #15140122). 1.5×10^5^ cells were seeded per well of Matrigel-coated 6-well plates (Corning, #3516). Cells were cultured for 30 days for astrocyte maturation, and the medium was changed every second day. Cells were passaged before reaching 80-90% confluency once per week. Following the initial 30-day differentiation period, astrocytes were maintained in serum-free astrocytes-medium. Before stimulation, the astrocyte medium was changed gradually to N2-medium over four days.

#### Generating microglia-assembled retinal organoids (iMG-RO)

3D-retinal organoids (3D-RO) were individually placed into 1.5 mL tubes (Roth, #1KP0.1), each containing 500 µL medium (either supplemented 3:1-medium or N2-medium depending on the age of the differentiation). preMG were collected as described in “Harvesting microglia precursor cells (preMG),” and 6×10^4^ cells were added to each organoid. After 72 hours, 6-8 organoids were pooled into one well of a 24-well plate (Corning, #3527) and cultured in the respective medium supplemented with 50 ng/mL MCSF for three weeks. Media was exchanged twice per week, and retinoic acid was added daily. Cultures were maintained at 37°C and 5% CO_2_ in a humidified incubator (BINDER C150).

### Generation of dissociated retinal organoid cultures

#### Retinal cup dissociation and 2D plating

At day 105, four 3D-retinal organoids were dissected to remove non-retinal tissue using a scalpel (Fisher Scientific, #11798343), transferred into a 1.5 mL tube, and washed twice in DPBS (Thermo Fisher Scientific, #14190250). Organoids were incubated in Accutase (STEMCELL Technologies, #07920) for 30 minutes at 37°C. Then, an equal volume of HBSS (Thermo Fisher Scientific, #14175-129) supplemented with 10% (v/v) heat-inactivated FBS (Thermo Fisher Scientific #10270106) was added, and organoids were dissociated by pipetting up and down ten times using a 200 µL tip. Cells were centrifuged at 3.2×g (VWR, Micro Star 17) for 2 minutes, the medium aspirated, and cells resuspended in N2-medium. Following another centrifugation at 3.2×g for 2 minutes, cells were resuspended in N2-medium supplemented with 20 ng/mL BDNF (Brain-Derived Neurotrophic Factor, Biolegend, #788904), passed through a 70 µm cell strainer (Corning, #352350), and distributed in 6 wells of a Matrigel-coated 8-well chamber (IBIDI, #80826). N2-medium supplemented with 20 ng/mL BDNF was changed every 3-4 days. 0.5 µM retinoic acid was added daily.

#### Integrating microglia precursor cells into dissociated retinal organoids

At day 130, preMG were harvested as described in “Harvesting microglia precursor cells (preMG),” and 6×10^4^ cells were added per well of an 8-well chamber. Cultures with and without microglia were maintained in N2-medium supplemented with 20 ng/mL BDNF and 50 ng/mL MCSF. Medium was exchanged every 3-4 days, and 0.5 µM retinoic acid was added daily.

| Plate | Surface area | Retinal cups/ well | preMG/ well |
| --- | --- | --- | --- |
| MEA | 0.5 cm<sup>2</sup> | 0.45 | 30.000 |
| 8 well IBIDI | 1 cm<sup>2</sup> | 0.75 | 60.000 |
| 24 well | 1.9 cm<sup>2</sup> | 1.5 | 125.000 |
| 12 well | 3.5 cm<sup>2</sup> | 3 | 250.000 |

### Stimulation paradigms

#### Stimulating microglia precursor cells to assay gene expression

preMG were harvested as described “Harvesting microglia precursor cells (preMG),” and 2×10^5^ cells were transferred into one well of a 24-well (Corning, #3527) containing 3:1-medium supplemented with 50 ng/mL MCSF. After 24h, the medium was replaced, and preMG were treated with poly(I:C) (Tocris, #4287) at a final concentration of 50 µg/mL. Untreated controls received 3:1-medium supplemented with 50 ng/mL MCSF. preMG were incubated for 24 hours at 37°C and 5% CO_2_. Four distinct cultures of independent differentiations were analyzed per condition.

#### Stimulating differentiated astrocytes

80% confluent astrocyte cultures were stimulated with poly(I:C) (Tocris, #4287) at a final concentration of 50 µg/mL for 24 hours. Untreated controls received N2-medium. Three distinct cultures of independent differentiations were analyzed per condition.

#### Stimulating microglia assembled dissociated retinal organoids

At week 20 (D139), N2-medium supplemented with 50 ng/mL MCSF was changed. We omitted the BDNF application because it has been shown to have anti-inflammatory effects ^17,18^. The withdrawal did not significantly alter ganglion cell survival (data not shown).

For each condition, one well of _diss_RO and iMG-_diss_RO were treated for 24 hours with either poly(I:C) (Tocris, #4287) at final concentration of 50 µg/mL only, 400 µM (S)-ibuprofen only, poly(I:C) plus (S)-ibuprofen in combination or poly(I:C) plus 20 nM SC-560 (Abcam, # ab120649) in combination. S-ibuprofen is the active enantiomere [152]. Untreated controls received medium supplemented with 50 ng/mL MCSF. Five distinct cultures of independent differentiations were analyzed per condition.

#### Stimulating retinal organoids

At week 20 (D139), three 3D-retinal organoids were transferred into one well of a 24-well plate per condition and cultured in N2-medium supplemented with 50 ng/mL MCSF. ROs in parallel to iMG-ROs were treated with either only poly(I:C) (Tocris, #4287) at a final concentration of 50 µg/mL or poly(I:C) plus 400 µM (S)-ibuprofen (Sigma-Aldrich, #375160-1G) in combination. Untreated controls received N2-medium supplemented with 50 ng/mL MCSF. Organoids were incubated for 24 hours at 37°C and 5% CO_2_. Five distinct cultures of independent differentiations were analyzed per condition.

#### CCL2 stimulation

At week 20 (D139), N2-medium supplemented with 50 ng/mL MCSF was changed, and BDNF was withdrawn from the medium. For each condition, one well of _diss_RO and iMG-_diss_RO were treated with recombinant human CCL2 (C-C Motif Chemokine Ligand 2, Peprotech, #300-04) at a final concentration of 10 ng/mL, 20 ng/mL, or 50 ng/mL. Untreated controls received N2-medium supplemented with 50 ng/mL MCSF. Five distinct cultures of independent differentiations were analyzed per condition.

### Gene expression profile of inflammatory cytokines

#### RNA isolation

Samples were washed with DPBS (Thermo Fisher Scientific, #14190250) before RNA isolation using the innuPREP RNA Mini Kit 2.0 (Analytik-Jena, #845-KS-2040050) as described in the manufacturer’s instructions. cDNA synthesis was performed with LunaScript RT SuperMix Kit (New England Biolabs, #E3010L) with a total RNA amount of 200-800 ng (same amount for each condition within experimental repetition) and stored at −20°C.

#### Gene expression analysis

RT-qPCR (Luna Universal qPCR Master Mix, New England BioLabs, #M3003L) was performed in 384-well plates (Bio-Rad; HSR4805) using the Roche Lightcycler 480 applying the device’s “Second Derivative Maximum Method.” The total reaction volume was 10 µL containing 1 µL of 1:10 diluted cDNA. The final concentration for each primer was 0.25 µM. The primer pairs are listed in **Supplementary Table 2**. Cycle conditions were 60 s at 95 °C for initial denaturation, followed by 40 cycles of denaturation (15 s; 95 °C) and annealing/extension (30 s; 60 °C). Each run was completed with a melting curve analysis to confirm the amplification of only one amplicon. Each PCR reaction was run in triplicates, from which a mean Cq value was calculated and used for further analysis. dCq values were obtained by normalizing mean Cq values to the geometric mean of four reference genes (GAPDH, ACTB, OAZ1, RPL27) measured within the same sample [equation 1]. ddCq values were then calculated by normalizing dCq values to the respective control condition (untreated cells/organoids) within each experimental repetition [equation 2]. Fold changes were obtained by transforming ddCq values from log2 to linear scale [equation 3].

Equations for consecutive RT-qPCR normalization:

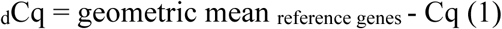

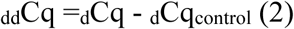

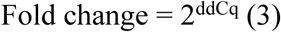

To analyze the mRNA expression relative to GAPDH, dCq values were normalized to GAPDH dCq values in the respective conditions.

### Proteome profiler array

Cultures were prepared as described in the sections “Retinal cup dissociation and 2D plating” and “Integrating microglia precursor cells into dissociated retinal organoids” with the following modification: Since samples were grown in 12-well plates, three retinal cups were dissociated per well, and 2.5×10^5^ preMG were added per well. Cultures were stimulated as described in “Stimulating microglia assembled dissociated retinal organoids”.

#### Human cytokine array

After two, four, or 24h of stimulation, the supernatant was harvested, snap-frozen on dry ice, and stored at −80°C. The Proteome Profiler Human Cytokine Array Kit (R&D Systems, #ARY005B) was performed following the manufacturer’s instructions. The membranes were imaged using the luminescent image analyzer Amersham Imager 600 (GE Healthcare Bio-Science). For the 24-hour time point, the supernatant from three distinct cultures of independent differentiations was assayed. Only one supernatant from one differentiation was screened for the two and four-hour time points.

#### Analysis of proteome profiler array

Pixel densities for positive signals were extracted using the ImageJ plugin ‘Protein Array Analyzer’ (https://imagej.nih.gov/ij/macros/toolsets/Protein%20Array%20Analyzer.txt). For each experimental condition, the average signals were determined per protein-of-interest, background signal subtracted, and signals normalized to the mean of six reference spots per membrane.

Hierarchical clustering of the median (24h time point) of normalized pixel values was carried out using the *pheatmap* package (version 1.0.12, RRID: SCR_016418) in R (version version 4.2.2). Fold changes were obtained by normalizing relative pixel values to the control condition.

### ELISA

Cultures were stimulated as described in “Stimulating microglia assembled dissociated retinal organoids.” After 24 hours of stimulation, the supernatant was harvested, snap-frozen on dry ice, and stored at −80°C. PGE2 ELISA (Enzo Life Sciences, #ADI-900-001) was performed according to the manufacturer’s instructions. Samples were analyzed in duplicates, and PGE2 concentration was determined based on the standard curve. The supernatant from three distinct cultures of independent differentiations was assayed.

### Histology

#### Fixation of 3D-retinal organoids

3D-retinal organoids were fixed in 4% (w/v) PFA (Paraformaldehyde, Thermo Fisher Scientific, #28908) in PBS for 25 minutes at room temperature on an orbital shaker in the dark. Then, organoids were washed three times with PBS at room temperature and cryopreserved in 30% (w/v) sucrose (Sigma-Aldrich, #84097) in PBS overnight at 4°C or stored in PBS at 4°C until further use.

#### Fixation of microglia precursor cells and dissociated retinal organoids

Cells were fixed in 4% (w/v) PFA in PBS for 20 minutes at room temperature in the dark, then washed three times with PBS at room temperature and stored in PBS at 4°C.

#### Cryostat sectioning

Cryopreserved 3D-retinal organoids were transferred to a cryomold (PolyScience, #18985) using a 1250 µL wide orifice pipette tip and embedded in Tissue-Tek O.C.T. compound (TTEK, A. Hartenstein) on dry ice. Samples were stored at −80°C until further use. Organoids were cut into 50 µm sections using a cryostat (MICROM, NX70 CRYOSTAR, Thermo Scientific). Sections were mounted onto glass slides Superfrost Plus (Lactan, #H867.1), dried at room temperature overnight, and stored at −80°C until further use. For immunostainings, slides were thawed and dried for 1 hour at room temperature. Sections on glass slices were encircled with an engraving, hydrophobic pen (Sigma-Aldrich, #Z225568).

#### Immunostaining of cryostat sections, microglia precursor cells, and dissociated retinal organoid cultures

Samples were incubated in a “blocking solution” containing 1% (w/v) bovine serum albumin (Sigma, #A9418), 5% (v/v) Triton X-100 (Sigma, #T8787), 0.5% (w/v) sodium azide (VWR, #786-299), and 10% (v/v) serum (either goat, Millipore, #S26, or donkey, Millipore, #S30) for two hours in a humidified chamber protected from light at room temperature. Afterward, the samples were immunostained with primary antibodies diluted in antibody solution containing 1% (w/v) bovine serum albumin, 5% (v/v) triton X-100, 0.5% (v/v) sodium azide, 3% (v/v) goat or donkey serum. They incubated overnight in a humidified chamber at room temperature. For the list of primary antibodies, see **Supplementary Table 3.** After washing the samples three times with PBS, the samples were incubated light-protected in a humidified chamber for 2 hours at room temperature, with the secondary antibodies diluted in antibody solution. The secondary antibodies raised in goat or donkey were purchased from Thermo Fisher Scientific (Alexa Fluor 488, Alexa Fluor 568, Alexa Fluor 647, 1:2000). The sections were washed three times with PBS. The nuclei were labeled with Hoechst 33342 (Thermo Fisher Scientific, Cat#H3570, 1:5000 diluted in PBS) for 15 minutes and then washed two times in PBS. After immunostaining, antifade solution [10% (v/v) mowiol (Sigma, #81381), 26% (v/v) glycerol (Sigma, #G7757), 0.2M tris buffer pH 8, 2.5% (w/v) Dabco (Sigma, #D27802)] was dropped on the cryostat sections and covered with microscope coverslips (Menzel-Glaser #0). Slides were dried at room temperature overnight. 8-well chambers were maintained in PBS. Samples were kept at 4°C for long-term storage.

#### Immunostaining of entire 3D-retinal organoids

The staining was performed as described under “*Immunostaining of cryostat sections, microglia precursor cells and dissociated retinal organoid cultures*” with the following adaptations: 3D-organoids were incubated in blocking for 24 hours on an orbital shaker at 4°C in the dark. The primary antibody concentration was doubled, and organoids were incubated for ten days on an orbital shaker at 4°C in the dark. After washing the organoids three times in PBS for 30 minutes each, secondary antibodies (Thermo Fisher Scientific, Alexa Fluor 488, and Alexa Fluor 647, 1:500) and Hoechst 33342 (1:1000) diluted in antibody solution were added for three days on an orbital shaker at 4°C in the dark. Finally, the organoids were washed three times in PBS for 30 minutes each. 4-5 3D-organoids were placed into one well of an 8-well chamber (IBIDI, #80826) and covered with 3% (w/v) low gelling temperature agarose (Sigma-Aldrich, #A9414-25G). The samples were stored in glycerol (Sigma-Aldrich, G7757) overnight at 4°C in the dark.

### Imaging and image analysis

#### Brightfield

The differentiation was monitored using a bright-field microscope (Olympus CKX41) with 5×, 10× and 20× objectives (Olympus) and a lens marker (Nikon), and an EVOS imaging system (Thermo Fisher Scientific) with 2×, 4×, 10×, 20×, 40× objectives (Thermo Fisher Scientific).

### Confocal microscopy

Images were acquired with a Zeiss LSM880 Airyscan or LSM800 inverted. For overview images, Plan-Apochromat 10× air objective NA 0.45 (WD=2.1mm) or Plan-Apochromat 20× Air objective NA 0.8 were used, and z-stacks were acquired. For detailed images, Plan-Apochromat 40× oil immersion objective NA 1.3 was used.

#### Imaging of dissociated retinal organoids

Three regions-of-interests were acquired per condition and biological replicate using the Plan-Apochromat 20× Air objective NA 0.8 with a zoom of 0.8.

#### Imaging of 3D retinal organoid sections

Based on the nuclei staining, one cryostat section per 3D retinal organoid displaying a retinal cup with a lumen was identified and imaged using the Plan-Apochromat 20×Air objective per condition and biological replicate.

#### Imaging of entire 3D-retinal organoids

The embedded organoids were imaged using Plan-Apochromat 10×Air objective NA 0.45 (WD=2.1mm).

#### Live cell imaging of dissociated microglia assembled retinal organoids

Microglia-assembled dissociated retinal organoids were generated as described in “Incorporating microglia-like cells into dissociated retinal organoids”. The cells were stimulated as described in “Stimulating microglia assembled dissociated retinal organoids”. Images were acquired with a Zeiss LSM880 inverted microscope and a Plan-Apochromat 20×/NA 0.8 Air objective in a temperature-controlled chamber (37°C). Z-stacked images of the Alexa 568 channel were captured every minute for 20 minutes.

#### Image analysis

Confocal images were converted to .ims files using the Imaris File Converter 9.9.1 and imported to Imaris 9.9 (Bitplane Imaris 3/4D Image Visualization and Analysis Software). Images were cropped and processed using background subtraction.

#### iMG positioning within layers

Cryostat sections with a focus on retinal cups were used for the analysis. Since we focused on retinal cups displaying a laminated structure, Hoechst-staining was used to identify the formation of layers. Microglia positioning was based on the location of the microglial cell soma. Each data point represents the percentage of microglia within a respective layer relative to the total number per section.

#### Determining the number of microglia in entire 3D-retinal organoids

Z-stack images of entire organoids were cropped to focus on the retinal cup. The number of microglia-like cells (iMG) were determined using the spot function of Imaris. The estimated XY diameter was set to 15 µm.

#### Determining cell numbers

The spot function of Imaris was used to analyze cells of interest and Hoechst^+^-cells. For nuclear stainings such as Hoechst, OTX2, BRN3, and KI67, the estimated XY diameter was set to 7 µm. For CALB2, CALB1, and RCVRN-labeling, the estimated XY diameter was set to 10 µm. To analyze the number of tdTomato^+^-iMG, the estimated XY diameter was set to 15 µm. The spots were manually edited. For cryostat sections of retinal organoids, we focused on the retinal cup.

For analysis, each data point represents the percentage of the respective marker relative to the total number of Hoechst^+^-cells, or the total number of microglia was determined for each region of interest. To calculate the proliferation rate of retinal cell types, the number of KI67^+^/iMG was subtracted from the total number of KI67^+^-cells, and the number of iMG were subtracted from the total number of Hoechst^+^-cells.

Fold change was determined by normalizing the median of three regions of interest per biological replicate to the respective control condition (untreated cells).

#### Determining Hoechst^+^-fragments

Hoechst^+^-fragments with a 2-5 µm diameter were manually counted using Imaris software. In the plot, each data point represents the number of Hoechst^+^-fragments per mm^2^.

#### Microglia surveillance

Time-lapse videos were binarized in ImageJ using the method ‘Li.’ The Matlab script determined the surveillance index [153], normalized to the total number of microglia imaged per video. Fold changes were calculated by normalizing to untreated control within each experimental repetition.

#### Microglia area

In Imaris, the surface rendering module with the surface detail set to 0.2 µm generated the microglia surface. iMG at the border of the image were removed. We excluded surfaces if multiple microglia were recognized as one surface. The exported Imaris file shows the surface area for each detected surface. In the plot, each data point represents the surface area of an individual microglia.

#### Microglia engulfing BRN3^+^-cells

Surface rendering was performed for iMG and BRN3^+^-cells with the surface detail set to 0.2 µm. The surface-surface co-localization function in Imaris was used to visualize co-localization. The total number of iMG and the number of iMG engulfing BRN3^+^-cells were determined using the spot function. In the plot, each data point represents the percentage of iMG engulfing a BRN3^+^-cell relative to the total number of microglia per field of view.

#### Distance from spot to surface

First, all BRN3^+^-cells were determined using the spot function of Imaris (XY diameter = 7 µm). Second, the iMG surface was generated using the surface rendering module with the surface detail set to 0.2µm. Finally, the function ‘spot to surface’ with a distance of 5 µm was used to determine the number of BRN3^+^-cells close to the iMG surface. In the plot, each data point represents the percentage of BRN3^+^-cells close to iMG relative to the total number of BRN3^+^-cells per field of view.

#### Graphics

All graphics were generated using R (version 4.2.2). Excel files were loaded into R via the *xlsx* package (version 0.6.5) [154]. Plots were made using ggplot2 (version 3.4.1) [155].

### Calcium imaging of dissociated cultures

#### AAV production and titration

The ISTA Molecular Biology Facility generated the virus. Human embryonic kidney (HEK) 293T cells were maintained at 37°C in 5% (v/v) CO_2_ in complete medium (DMEM medium (Thermo Fisher Scientific, #31966047), 10% (v/v) fetal bovine serum (Thermo Fisher Scientific, #10270106], 1% (v/v) penicillin/streptomycin (Thermo Fisher Scientific, #15140122), 1% (v/v) non-essential amino acids (Sigma-Aldrich, #M7145-100ML). Ten 15-cm culture dishes with 80% confluency were transfected using 6.8 μM polyethyleneimine (Polysciences, #24765-1), 70 µg AAV transgene plasmid (pAAV-EF1a-GCaMP6s-WPRE-pGHpA, Addgene, #67526), 70 µg 7M8 Cap-encoding plasmid (7M8, Addgene, #64839), 200 µg pHGT1-Adeno1 helper plasmid. Sixty hours post-transfection, cells were harvested with a cell scraper and pelleted at 4000 rpm for 15 minutes at 4°C. The pellet was resuspended in lysis buffer (200 mM NaCl, PBS, 0.001% pluronic F68, sterile filtered). Cell-lysis was obtained by three rounds of freezing-thawing cycles between dry ice/ethanol and a 37°C water bath, followed by sonication for 1 minute at 37kHz. Next, Benzonase (50 U/mL, Sigma Aldrich, #E1014-25KU) was added, and the solution was incubated at 37°C for 45 minutes. Afterward, the solution was centrifuged at 2415×g for 10 minutes at 4°C. The AAV particles in the supernatant were purified by discontinuous iodixanol gradient ultracentrifugation. Optiseal tubes (Beckman Coulter, 361625) were filled with a density gradient of 60%, 40%, 25%, and 17% iodixanol solutions (Optiprep Iodixanol, Progen Biotechnik, 1114542). The virus lysate was loaded on the top layer, and the tubes were centrifuged at 350000 g (Beckman Optima XPN-100 ultracentrifuge Sorvall T-850 rotor) for 90 minutes at 10°C. The AAV particles were harvested from the intersection of 60% and 40% gradients and concentrated using 100kDa Vivaspin 20 Centrifugal Concentrator. Aliquots were stored at −80°C.

For titration by qPCR, AAV particles were denatured with DNase I (Fisher Scientific, #10103533) and the viral DNA quantification was performed with Universal SYBR Master Mix 2X (Thermo Fisher Scientific, #4309155) using the following primers: forward primer: 5’- GGAACCCCTAGTGATGGAGTT; reverse primer: 5’-CGGCCTCAGTGAGCGA. The final titer measured 1.1×10^13^ viral genome copy number per milliliter (GC/mL).

#### AAV infection of dissociated cultures

At week 17 (D120), dissociated retinal organoid cultures were infected with 5×10^10^ GC of AAV2/7m8-EF1α-GCAMP6s [10, 110] cultured in 100 µL 3:1-medium. After 24 hours, 100 µL fresh 3:1-medium was added. The next day, the medium was changed to N2-medium supplemented with 20 ng/mL BDNF. The medium was changed every 3 to 4 days, and 0.5 µM retinoic acid was added daily.

#### Calcium imaging

Four days before calcium imaging at day 138, dissociated retinal organoids were gradually transferred to BrainPhys medium (StemCell Technologies, #05791) supplemented with 1×N2 supplement, 100 μM taurine supplemented with 20 ng/mL BDNF, 50 ng/mL MCSF and 0.5 µM retinoic acid. Twenty-four hours before imaging, the medium was changed, and BDNF was withdrawn. Cultures were treated as described in “Stimulating microglia assembled dissociated retinal organoids”, except that the samples were cultured in a supplemented BrainPhys medium. Live imaging was performed using the Dragonfly microscope (Andor Dragonfly 505, Oxford Instruments) equipped with a heated chamber at 37°C and CFI P Apochromat 20× NA 0.95/ WD 0.95 mm water objective (Nikon, MRD77200). The Andor iXon Ultra 888Ultra EMCCD camera (13 μm pixel size) was used to acquire the 488 nm channel using a pinhole size of 25 µm. The following parameters were used for acquisition: exposure time of 40 ms, EM gain of 100, Laser 7.0%, and an AOI of 1024×1024. Baseline activity was acquired for five minutes using a time series at 12.16 Hz. Baseline calcium dynamics were recorded for 5 minutes from five distinct cultures of independent differentiations.

#### Pharmacological manipulation

First, baseline calcium activity was recorded for 2.5 minutes. For pharmacological manipulation, either 1 µM Tetrodotoxin (Abcam, # ab120054) to block voltage-gated sodium channels was applied or a mixture of 10 μM NBQX (2,3-dioxo-6-nitro-7-sulfamoyl-benzo[f]quinoxaline; Tocris Bioscience, #1044), 10 μM DL-AP4 (DL-2-amino-4-phosphonobutyric acid, Tocris Bioscience, #0101), 10 μM (±)-CPP (3-[(R)-2-carboxypiperazin-4-yl]-propyl-1-phosphonic acid; Abcam, #ab144495) to inhibit glutamatergic synaptic transmission was applied and. After 5 minutes of incubation, calcium activity was recorded for another 2.5 minutes.

To chelate extracellular calcium, 5mM EGTA (ethylene glycol-bis(β-aminoethyl ether)-N,N,N′,N′-tetraacetic acid) was applied and recording immediately continued for another 2 minutes.

#### Calcium imaging analysis

Cells showing calcium transients were identified manually as regions of interest using ImageJ, and mean grey values were extracted. We focused on cells where the cell soma and the primary branches were clearly visible. Transients in the background were not included in the analysis. Fluorescent signal time series (F/ΔF: change in fluorescence divided by the mean baseline fluorescence) were calculated for each region of interest. Calcium events were detected in Matlab 2017 using the script ‘PeakCaller’ [156] using the following parameters: required rise = 9% absolute; max. lookback =100 pts; required fall = 5% absolute; max. lookahead = 100 pts; no trend control; trend smoothness = 100; interpolate across closed shutters = true. For each cell, waveform parameters such as number of events and peak amplitude were extracted and plotted.

### Statistical analysis

All statistics were performed using R (version 4.2.2). The linear regression model was calculated using the “lme4”-package. First, groups for comparison were tested for normal distribution and equal variances using the Shapiro-Wilk and Levene tests, respectively. No data was excluded for analysis. If the data was normally distributed and the groups had equal variances, Student’s t-test was used to compare the two groups. For multiple comparisons, the default contrast for unordered variables was set to ‘contr.sum’ to perform one-way ANOVA, followed by Tukey’s post-hoc comparison. Welch’s test was performed to compare normally distributed groups with unequal variances. A two-sided one-sample T-test was used to analyze if a normally distributed condition significantly differs from a value of 1 or 0.

If groups were not normally distributed, the Wilcoxon rank-sum test or Kruskal-Wallis test, followed by Dunn’s test, was used to compare two or multiple groups, respectively. For multiple comparisons, p-values were adjusted using the “p.adjust” function, and the method was set to ‘BH.’ The following packages were used to perform the analysis: “FSA”-package (dunn-test); “multcomp”-package (Tukey’-test); “psych”-package (t-test, kruskal-wallis test); “stats”-package (wilcox-test, shapiro-wilk test), “dplyr”-package (levene-test).

Significance levels are indicated using the following notation: ^n.s.^p > 0.05; ^∗^p < 0.05; ^∗∗^p < 0.01; ^∗∗∗^p < 0.001. Details about the statistical analysis are summarized in **Supplementary Table 4**, and the respective raw data in **Supplementary Table 5**.

---

**Supplementary Figure 1-.**
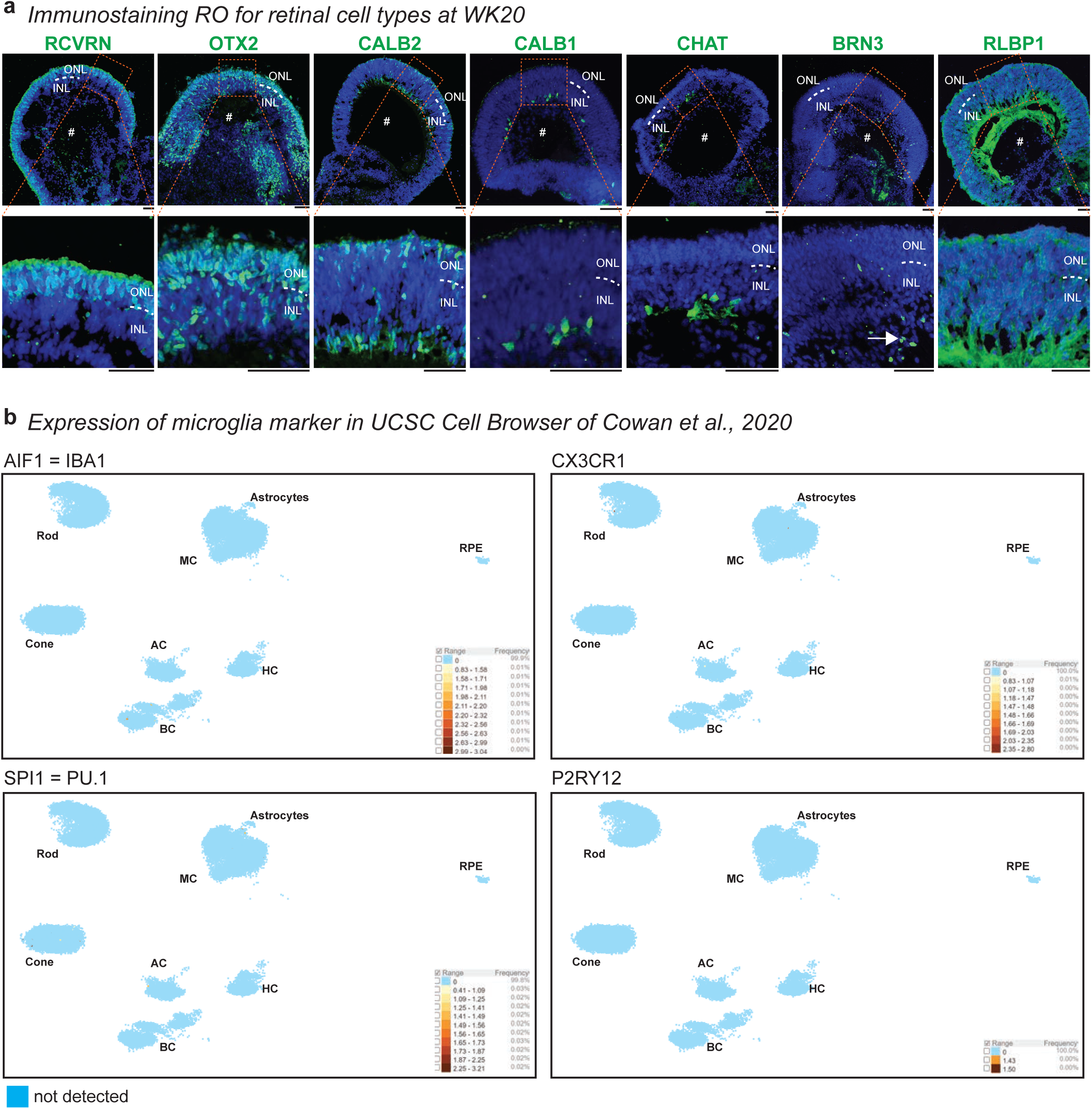
Characterization of 3D retinal organoids. **a**, Representative cryostat section images of 3D-retinal organoid counterstained with the nuclei-dye Hoechst (blue) and immunostained for retinal cell type-specific markers (green) and at week 20: RCVRN (recoverin; photoreceptors). OTX2 (orthodenticle homeobox 2; photoreceptors, bipolar cells). CALB2 (calretinin; photoreceptors, bipolar-, amacrine cells). CALB1 (calbindin; amacrine-, horizontal cells). CHAT (choline acetyltransferase; amacrine cells). BRN3 (brain-specific homeobox/POU domain protein 3B; ganglion cells). RLBP1 (cellular retinaldehyde-binding protein; Müller glia). ONL: outer nuclear layer. INL: inner nuclear layer. White dashed line: outer plexiform layer. #: retinal cup lumen. White arrow: BRN3^+^-cells close to lumen. Scale bar: 50 µm. **b**, Expression of microglia transcript markers in USCS Cell Browser of Cowan *et al*., 2020: Dataset ID: ‘Developed human retinal organoid.’ Uniform manifold approximation and projection (UMAP) of transcript expression for AIF (also known as IBA1, ionized calcium-binding adapter molecule 1), CX3CR1 (C-X3-C motif chemokine receptor 1), SPI1 (also known as PU.1, Spi-1 proto-oncogene) and P2RY12 (purinergic receptor P2Y12) of 3D-retinal organoid at week 32 and 38. AC: amacrine cell. BC: bipolar cell. Cone: cone photoreceptors. HC: horizontal cell. MC: Müller glia. RPE: retinal pigment epithelium. Rod: rod photoreceptors. Blue dot: not detected.

**Supplementary Figure 2-.**
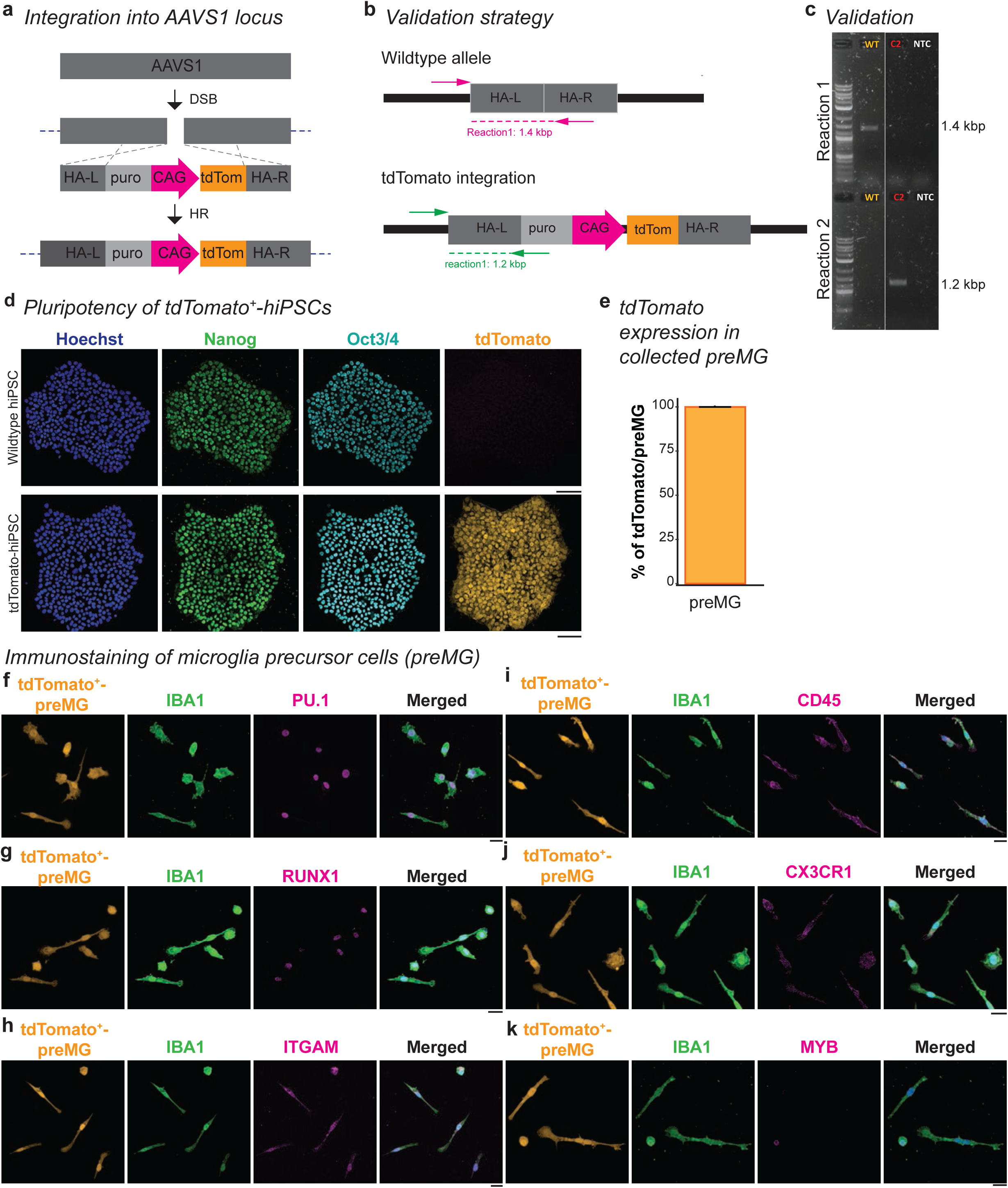
Generation of tdTomato^+^-hIPSC cell line and characterization of differentiated tdTomato^+^-microglia precursor cells (preMG). ***a****, Integration strategy into the* adeno-associated virus integration site 1 (*AAVS1) locus.* DSB: double-strand break. CAG: CMV immediate enhancer/β-actin promoter. HA-L: homologous arm left. HA-R: homologous arm right. HR: homologous recombination. *Puro: puromycin selection side. tdTom: tdTomato*. ***b****-**c**, Validation strategy. Reaction 1: wildtype allele: PCR product 1.4 kbp. Reaction 2: tdTomato allele: PCR product 1.2 kbp.* PCR: polymerase chain reaction. ***c****, PCR product size*. Top: Reaction 1 - wildtype AAVS1 locus (1.4 kbp). Bottom: Reaction 2 - construct integrated into AAVS1 (1.2 kbp). Orange: wildtype clone. Red: clone with homozygous integration of the construct. NTC: non-template control. Kbp: kilobase pair. **d**, Validating pluripotency for the wildtype human induced pluripotent stem cell (hIPSC) line SC102A (top) and the tdTomato^+^-hiPSC line SC102A (bottom). Immunostaining of hIPSC colonies for NANOG (nanog homeobox, green), OCT3/4 (octamer-binding protein 3, cyan), and counterstaining for the nuclei-dye Hoechst (blue). Intrinsic tdTomato expression (orange). Scale bar: 100 µm. **e**, Bar chart of tdTomato^+^/ IBA1^+^-preMG with standard error of the mean. **f-k**, Representative images of tdTomato-expressing microglia precursor cells (preMG, orange) harvested from the supernatant and plated on a new dish. Cells counterstained for the nuclei-dye Hoechst (blue, merged image), immunostained for IBA1 (ionized calcium-binding adapter molecule 1, green) and the microglia/macrophage markers in magenta for **f**, PU.1 (hematopoietic transcription factor PU.1); **g**, RUNX1 (runt-related transcription factor 1); **h**, ITGAM (integrin subunit alpha m); **i**, CD45 (cluster of differentiation 45/ protein tyrosine phosphatase receptor); **j**, CX3CR1 (chemokine (C-X3-C) receptor 1); **k**, MYB (MYB proto-oncogene). Scale bar: 20µm.

**Supplementary Figure 3-.**
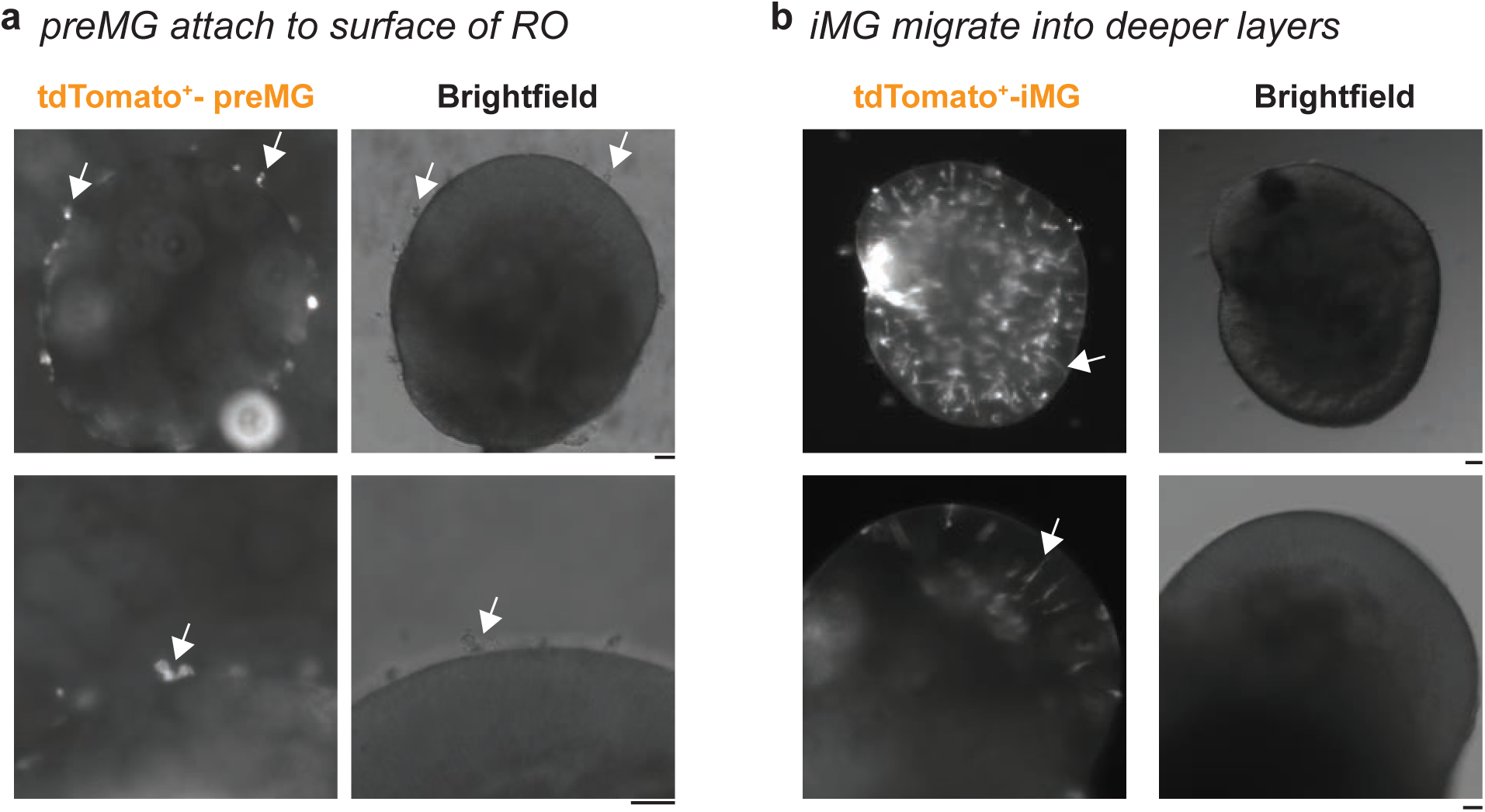
tdTomato^+^-microglia precursor cells (preMG) integration patterns into 3D retinal organoids. Representative images of 3D-retinal organoids. Left: fluorescence image, right: brightfield image. 4x magnification (top) and 10x magnification (bottom). Scale bar: 20µm. **a**, tdTomato^+^-microglia precursor cells (preMG, white arrow) attach on week 17 at the surface of 3D-retinal organoids (RO). **b**, tdTomato^+^-microglia-like cells (iMG) integrate into the 3D-retinal organoids at week 20, showing a bipolar shape (white arrow).

**Supplementary Figure 4-.**
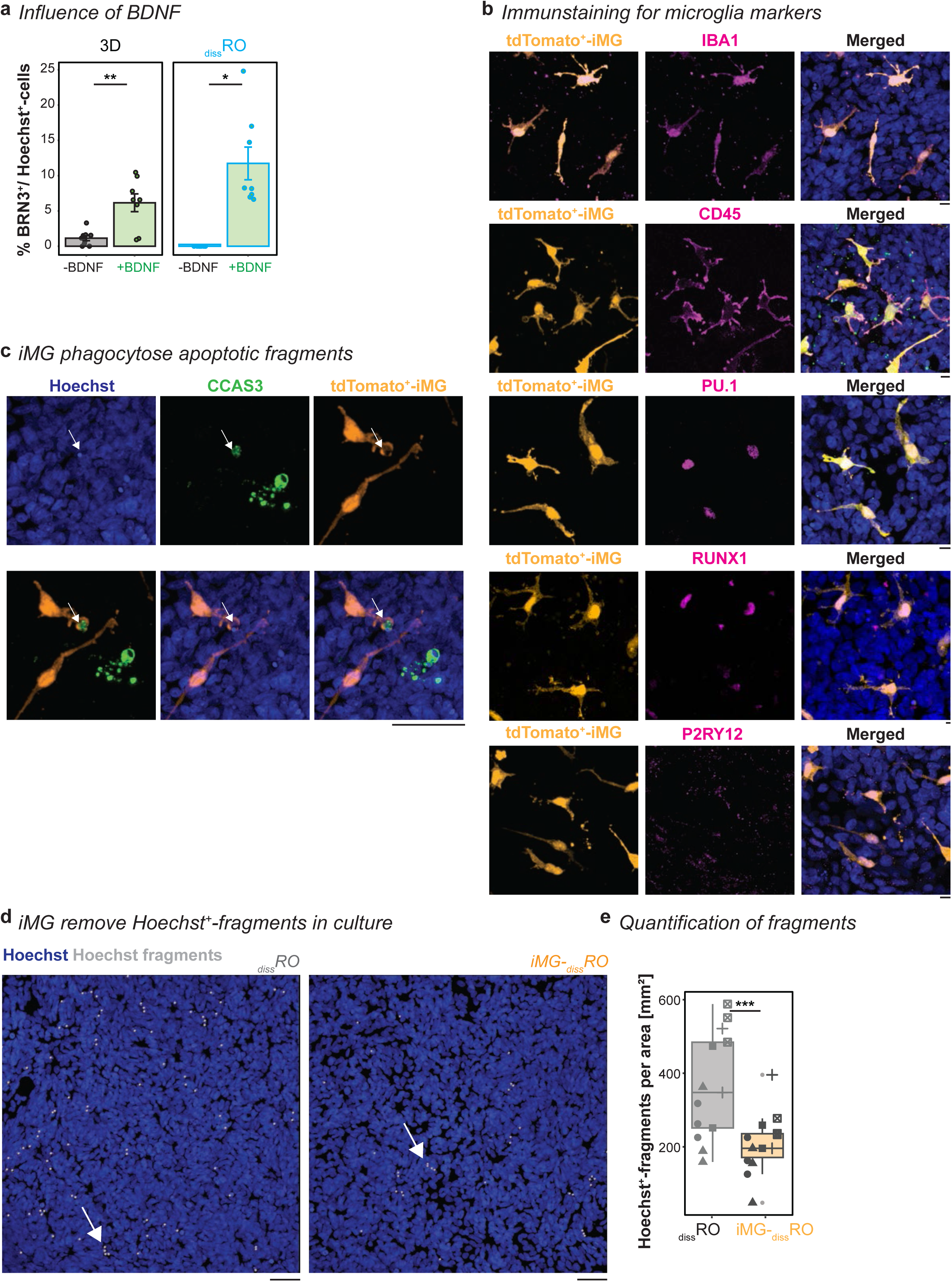
Validate iMG signature and their consequences in the dissociated retinal organoid model. **a**, Impact of brain-derived neurotrophic factor (BDNF) on ganglion cell survival. Bar chart of percentage of BRN3^+^-cells relative to Hoechst^+^-cells with standard error of the mean in 3D-retinal organoids (RO, left) and dissociated retinal organoid culture (_diss_RO, right) cultured either in standard retinal organoid differentiation media without (-BDNF, grey) or supplemented with BDNF (+BDNF, green) from week 15 to 20. RO: Each dot is one cryostat section of independent retinal cups. Welch’s t-test. _diss_RO: Each dot is one region of interest. One-sample Wilcoxon signed rank test. **b-d**, Representative images of iMG-_diss_RO for tdTomato expression (orange), counterstained for the nuclei-dye Hoechst (blue) and immunostained for **b**, macrophage/microglia-marker (magenta): IBA1 (ionized calcium-binding adapter molecule 1), CD45 (cluster of differentiation 45/ protein tyrosine phosphatase receptor), PU.1 (hematopoietic transcription factor PU.1), RUNX1 (runt-related transcription factor 1), and P2Y12 (purinergic receptor P2Y G-protein-coupled 12). Scale bar: 10 µm. **c**, the apoptotic marker CCAS3 (cleaved caspase3, green). White arrowhead: iMG engulfing CCAS3^+^-fragment. Scale bar: 50µm. **d**, Hoechst^+^-fragments (white) in _diss_RO (left) and iMG-_diss_RO (right). Scale bar: 50 µm. **e**, Boxplot of percent of Hoechst^+^-fragments per area in _diss_RO (grey) and iMG-_diss_RO (orange). Symbols: single ROI of three biological replicates from five independent differentiations. Welch’s t-test. For detailed statistical analysis, see **Supplementary Table 4**. ***p < 0.001. **p < 0.01. *p < 0.05.

**Supplementary Figure 5-.**
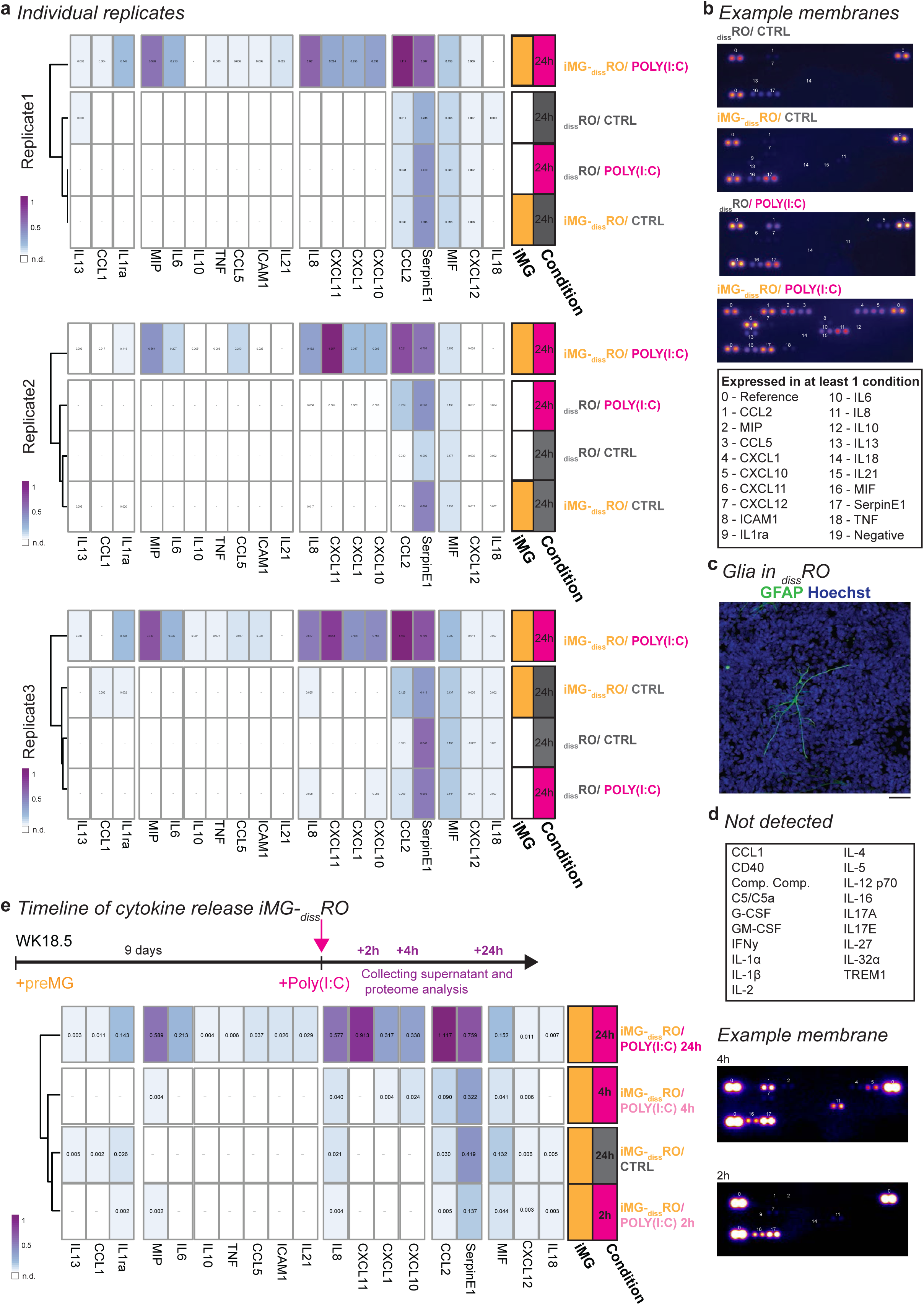
Individual inflammatory proteome profiler results. **a**, Release of inflammatory cytokines and chemokines into the supernatant based on the experimental paradigm described in Figure 3a for control (CTRL, grey) and 24h-poly(I:C) (magenta) stimulation. Individual heatmap plots with color-coded mean pixel intensity relative to the reference of three independent differentiations White: n.d. (not detectable). Side-bar: condition with iMG (orange) *versus* without (white) or CTRL *versus* poly(I:C). **b**, Representative membranes for each condition. Numbers refer to the legend below. **c**, Example images of _diss_RO counterstained with the nuclei-dye Hoechst (blue) and immunostained for the glial marker GFAP (glial fibrillary acidic protein, green). Scale bar: 20 µm. **d**, List of proteins assayed on the membrane but not detected in the supernatant of any condition. **e**, Same assay as for **a** with additional measurement of cytokine and chemokine release after two and four hours compared to 24h in iMG-_diss_RO with annotated example membranes iMG-_diss_RO.

**Supplementary Figure 6-.**
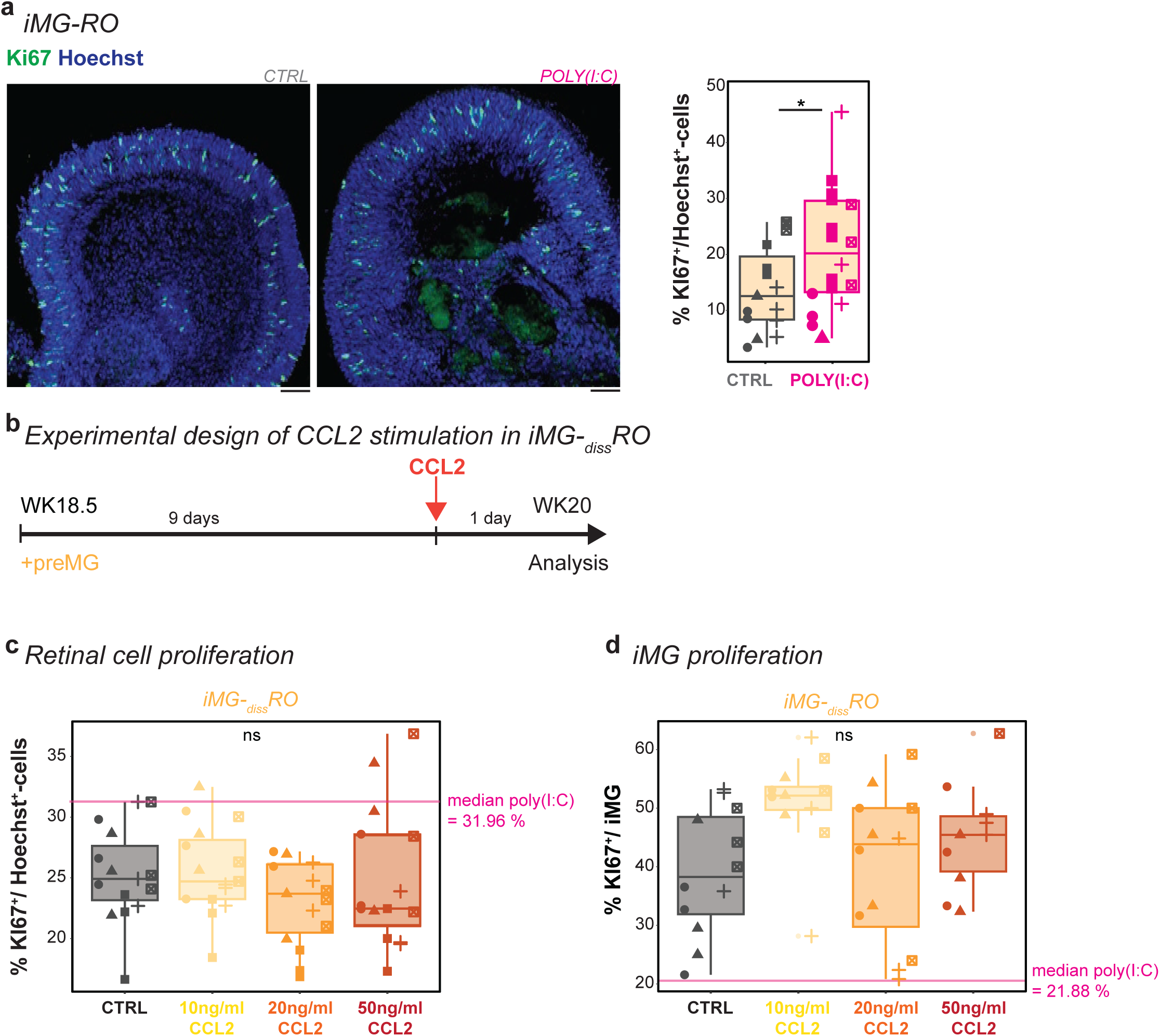
The poly(I:C)-mediated proliferation rate increase cannot be replicated with CCL2 alone. **a**, Representative image of cryostat section with a focus on the retinal cup of iMG-RO counterstained with the nuclei-dye Hoechst (blue) immunostained for the proliferation marker KI67 (green) after 24 hours of untreated control (CTRL, grey, left) or poly(I:C) stimulation (magenta right). Scale bar: 50 µm. Next: Boxplot of KI67^+^-cells relative to Hoechst^+^-cells per section excluding K67^+^/iMG. Each symbol: an independent differentiation (n=5). Student’s t-test. **b**, Experimental timeline. At WK18.5, preMG are added to _diss_RO. After nine days, cultures received fresh medium for control (CTRL, grey) or CCL2 stimulation iMG-_diss_RO for 24 hours. **c**, Effect of CCL2 on retinal cell proliferation excluding iMG. Boxplot of percent KI67^+^-cells relative to Hoechst^+^-cells in iMG-_diss_RO for CTRL and CCL2 stimulation at a final concentration of 10 ng/mL (yellow), 20 ng/mL (orange), and 50 ng/mL (red). Magenta line: Median proliferation rate in poly(I:C) stimulation of iMG-_diss_RO (Figure 4g). Symbols: three biological replicates from five independent differentiations. One-way ANOVA. **d**, Effect of CCL2 on iMG proliferation. Boxplot of percent KI67^+^/iMG for CTRL and CCL2 stimulation at a final concentration of 10 ng/mL (yellow), 20 ng/mL (orange), and 50 ng/mL (red). Magenta line: Median iMG-proliferation rate in poly(I:C) stimulation of iMG-_diss_RO (Figure 4d). Symbols: three biological replicates from five independent differentiations. Kruskal-Wallis test. For detailed statistical analysis, see **Supplementary Table 4**. *p < 0.05. ^ns^p > 0.05, not significant.

**Supplementary Figure 7-.**
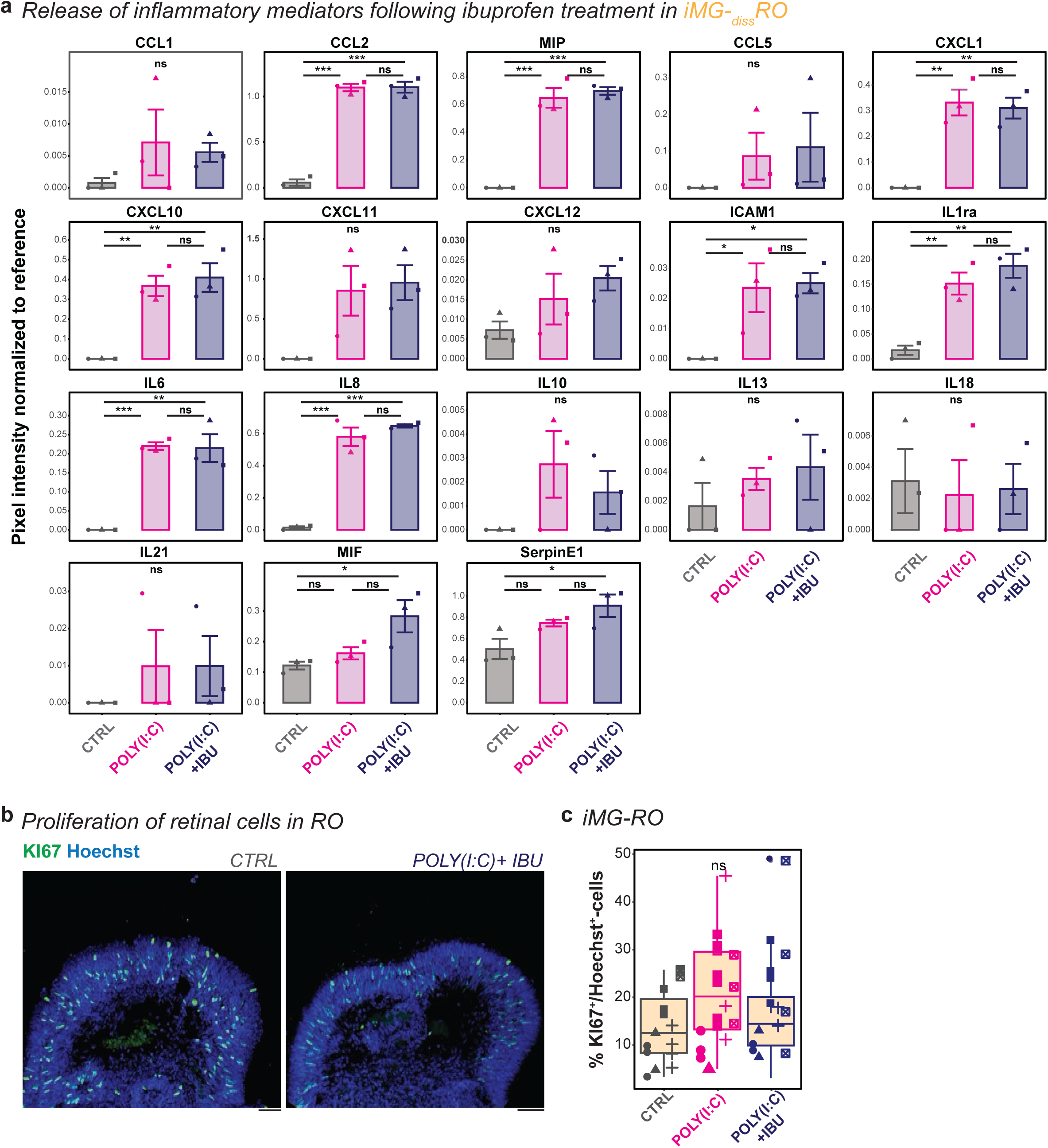
Comparison of individual secreted inflammatory mediators after ibuprofen exposure. **a**, Release of inflammatory cytokines and chemokines into the supernatant based on the experimental paradigm described in Figure 3a for control (CTRL, grey), poly(I:C) (magenta), and poly(I:C) and S(+)-ibuprofen (poly(I:C)+IBU, blue) stimulation. Release of different inflammatory mediators into the supernatant of iMG-_diss_RO. Bar chart of pixel intensity normalized to reference with standard error of the mean for CTRL, poly(I:C), and poly(I:C)+IBU. Each symbol: an independent differentiation (n=3). One-way ANOVA with post-hoc Tukey’s test except IL13, IL18, IL21 Kruskal-Wallis test. **b-c**, Ibuprofen treatment of iMG-RO. **b**, Representative images of cryostat sections focusing on retinal cup of iMG-RO for CTRL (left) and poly(I:C)+IBU (right) at WK20 counterstained for the nuclei-dye Hoechst (blue) and the proliferation marker KI67 (green). Scale bar: 50µm. **c**, Boxplot of percent of KI67^+^-cells relative to Hoechst^+^-cells excluding KI67^+^/iMG per section of an independent iMG-RO for CTRL, poly(I:C), and poly(I:C)+IBU. Each symbol: an independent differentiation (n=5). Kruskal-Wallis test. For detailed statistical analysis, see **Supplementary Table 4**. ***p < 0.001. **p < 0.01. *p < 0.05. ^ns^p > 0.05, not significant.

**Supplementary Figure 8-.**
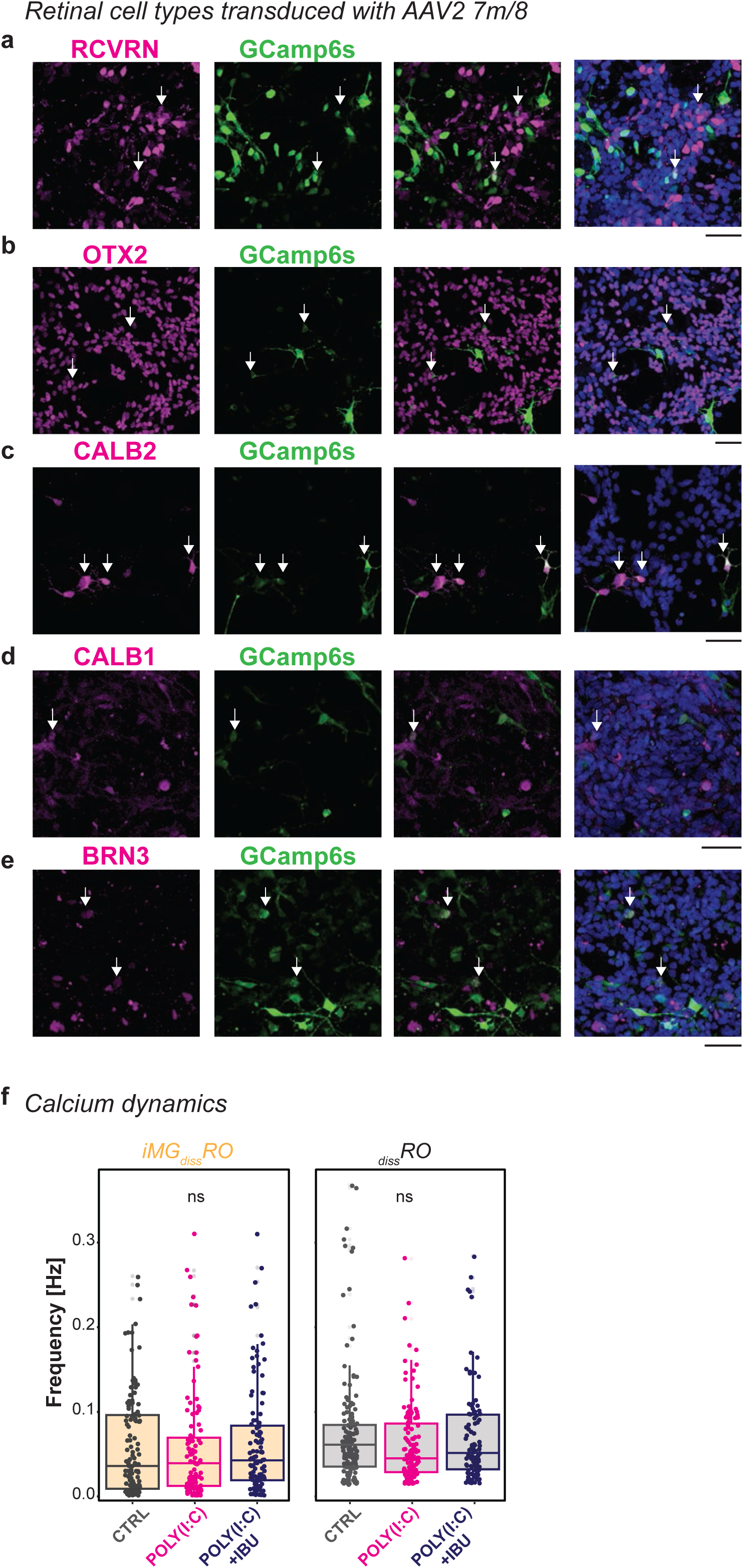
GCAMP6s expression across retinal cell types. **a-e**, Example ROI images of _diss_RO infected with AAV2-GCAMP6s at WK17, analyzed at WK20, counterstained for the nuclei-dye Hoechst (blue) and the calcium sensor GCAMP6s (green), and immunostaining for retinal cell types (magenta): **a**, RCVRN (recoverin; photoreceptors). **b**, OTX2 (orthodenticle homeobox 2; photoreceptors, bipolar cells). **c**, CALB2 (calretinin; photoreceptors, bipolar-, amacrine cells). **d**, CALB1 (calbindin; amacrine-, horizontal cells). **e**, BRN3 (brain-specific homeobox/POU domain protein 3B; ganglion cells). Arrow: Co-expression of calcium sensor and retinal marker. Scale bar: 50µm. **f**, Spontaneous calcium dynamics in iMG-_diss_RO (orange) and _diss_RO (grey) for control (CTRL, grey), poly(I:C) (magenta), and poly(I:C) and S(+)-ibuprofen (poly(I:C)+IBU, blue) stimulation. Boxplot of the mean frequency [Hz] during five minutes of recording. Each dot represents an active cell. Recordings from five biological replicates from independent differentiations. Kruskal-Wallis test. For detailed statistical analysis, see **Supplementary Table 4**. ^ns^p > 0.05, not significant.

**Supplementary Figure 9-.**
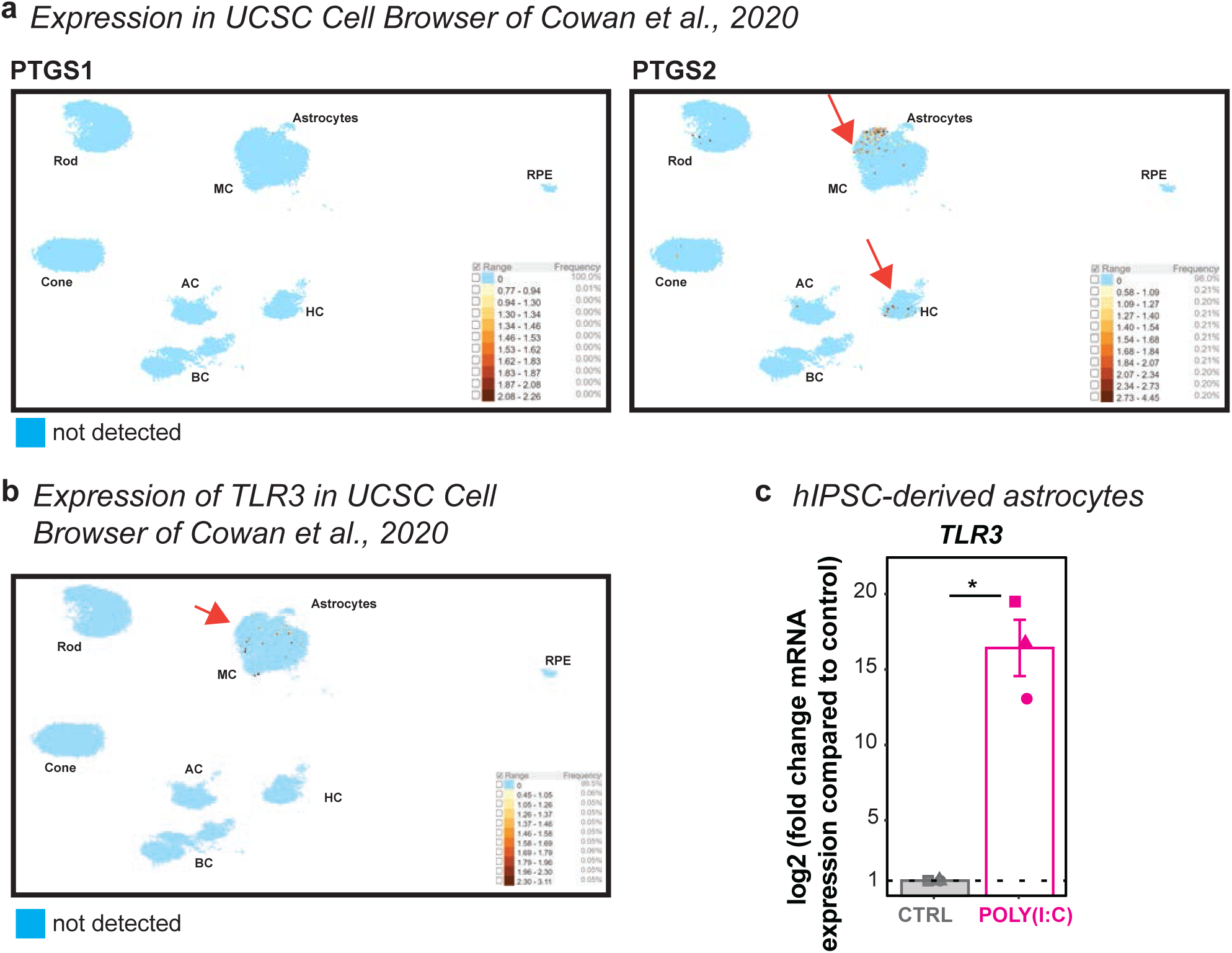
PTGS1, PTGS2, and TLR3 mRNA expression profile. **a-b**, Expression of (**a**) PTGS1 and PTGS2 (prostaglandin-endoperoxide synthase 1 and 2) as well as (**b**) TLR3 (toll-like receptor 3) in USCS Cell Browser of *Cowan et al., 2020*. Cell Browser dataset ID: ‘Developed human retinal organoid.’ Uniform manifold approximation and projection (UMAP) of transcript expression. AC: amacrine cell. BC: bipolar cell. Cone: cone photoreceptors. HC: horizontal cell. MC: Müller glia. RPE: retinal pigment epithelium. Rod: rod photoreceptors. Red arrow: positive transcript expression. Blue dot: not detected. **c**, Real-time quantitative polymerase chain reaction (RT-qPCR) for TLR3 (toll-like receptor 3) in hIPSC-derived astrocytes for untreated control (CTRL, grey) and poly(I:C) (magenta) exposure. Mean mRNA transcript log2-fold changes compared to untreated control cells with standard error of the mean. Each symbol is an independent differentiation (n=3). One sample t-test. For detailed statistical analysis, see **Supplementary Table 4**. *p < 0.05.

**Supplementary Video 1 – Microglia surveillance control.**

**Twenty** minutes of iMG (grey) motility recording after 24-hour exposure to fresh medium (control) in iMG-_diss_RO at WK20. One z-stack is imaged every minute. Maximum intensity projection. Scale bar: 50 µm.

**Supplementary Video 2 – Microglia surveillance poly(I:C).**

Twenty minutes of iMG (grey) motility recording after 24-hour exposure to poly(I:C) in iMG-_diss_RO at WK20. One z-stack is imaged every minute. Maximum intensity projection. Scale bar: 50 µm.

**Supplementary Video 3 – Microglia surveillance poly(I:C)+IBU.**

Twenty minutes recording of iMG (grey) motility after 24-hour exposure to poly(I:C) and S(+)-ibuprofen (poly(I:C)+IBU) exposure in iMG-_diss_RO at WK20. One z-stack is imaged every minute. Maximum intensity projection. Scale bar: 50 µm.

**Supplementary Video 4 – Spontaneous calcium dynamics.**

Three hundred seconds baseline recording of **a**, iMG-_diss_RO and **b**, _diss_RO infected with AAV2 7m8 EF1α GCAMP6s. Video shows cropped region of interest. Frames are indicated in the upper left corner. Scale bar: 10 µm.

**Supplementary Video 5 – TTX abolishes calcium transients.**

150 seconds baseline recording of _diss_RO (frame 0-1993). Five minutes incubation in TTX followed by another 150 seconds recording (frame 1994-3320). Video shows cropped region of interest. Frames are indicated in the upper left corner. Scale bar: 10 µm.

**Supplementary Video 6 – Glutamatergic modulation.**

150 seconds baseline recording of _diss_RO (frame 0-1696). Five minutes incubation with glutamatergic blockers CPP, NBQX, and APB followed by another 150-second recording (frame 1697-3320). Video shows cropped region of interest. Frames are indicated in the upper left corner. Scale bar: 10 µm.

**Supplementary Video 7 – Calcium chelator EGTA abolishes calcium transients.**

A 180-second baseline recording of iMG-_diss_RO (frame 0-1990) paused at frame 1990 to add EGTA, followed by another 120-second recording (frame 1991-3319). Video shows cropped region of interest. Frames are indicated in the upper left corner. Scale bar: 10 µm.

**Supplementary Table 1 – Overview of human induced pluripotent stem lines (hIPSC) included in this study.**

hPSCreg.eu, human pluripotent stem cell registry. MYC, MYC proto-oncogene. KLF4, kruppel-like factor 4. Large T antigen, large tumor antigen. LIN28, zinc finger CCHC domain-containing protein. OCT4 (octamer-Binding Protein 4)/ POU5F1 (POU domain, class 5, transcription factor 1). SOX2, sex-determining region Y-box 2. SV40, simian-virus 40.

**Supplementary Table 2 – List of antibodies.**

**Supplementary Table 3 – Primer sequences.**

**Supplementary Table 4 – Description of the statistical analysis.**

**Supplementary Table 5 – Raw data.**

## Acknowledgments

We thank the scientific service units at ISTA, specifically the Lab Support Facility (LSF), the Molecular Biology Services/Virus Services Team, specifically Flavia Gama Gomes Leite and Mark Andrew Smyth, for the virus production, and the Imaging and Optics Facility (IOF). We thank all members of the Siegert group and Marco Benevento for their constant feedback on the project and comments on the manuscript. A special thanks to Rouven Schulz for input on statistical analysis and sharing R-scripts, Gloria Colombo for the introduction to cell sorting, and Negar Vehdani for her support in cell culture. This research was supported by the Gesellschaft für Forschungsförderung Niederösterreich (grant No. Sc19-017 to V.H.).

## Author contributions

Conceptualization, V.S. and S.S.; investigation, validation, and methodology, V.S., M. K-D, A.V., S.S.; formal analysis, V.S.; writing original draft and visualization, V.S., S.S; writing – review & editing, V.S., M. K-D, A.V., S.S.; supervision, funding acquisition, S.S.

## Declaration of interest

The authors declare no competing interest.

## Notes

### Competing Interest Statement

The authors have declared no competing interest.

